# Barley cultivars shape the abundance, phenotype, genotype and gene expression of their associated microbiota by differential root exudate secretion

**DOI:** 10.1101/2023.07.05.547901

**Authors:** Alba Pacheco-Moreno, Anita Bollmann-Giolai, Govind Chandra, Paul Brett, Jack Davies, Owen Thornton, Philip Poole, Vinoy Ramachandran, James K.M. Brown, Paul Nicholson, Chris Ridout, Sarah DeVos, Jacob G. Malone

**Affiliations:** John Innes Centre, Norwich Research Park, Colney Lane, Norwich NR4 7UH; University of Zürich, Winterthurerstrasse 190, 8057 Zürich, Switzerland; IBERS, Aberystwyth University, Aberystwyth, Ceredigion, SY23 3EE; Rothamsted Research, Harpenden, AL5 2JQ; Department of Biology, University of Oxford, South Parks Road, Oxford, OX1 3RB; New Heritage Barley, Norwich Research Park, Norwich, NR4 7UG; School of Biological Sciences, University of East Anglia, Norwich NR4 7TJ

## Abstract

Plant associated microbes play vital roles in promoting plant growth and health, with plants secreting root exudates into the rhizosphere to attract beneficial microbes. Exudate composition defines the nature of microbial recruitment, with different plant species attracting distinct microbiota to enable optimal adaptation to the soil environment. To more closely examine the relationship between plant genotype and microbial recruitment, we analysed the rhizosphere microbiomes of landrace (Chevallier) and modern (NFC Tipple) barley cultivars. Distinct differences were observed between the plant-associated microbiomes of the two cultivars, with the plant-growth promoting rhizobacterial genus *Pseudomonas* substantially more abundant in the Tipple rhizosphere. Striking differences were also observed between the phenotypes of recruited *Pseudomonas* populations, alongside distinct genotypic clustering by cultivar. Cultivar-driven *Pseudomonas* selection was driven by root exudate composition, with the greater abundance of hexose sugars secreted from Tipple roots attracting microbes better adapted to growth on these metabolites, and vice versa. Cultivar-driven selection also operates at the molecular level, with both gene expression and the abundance of ecologically relevant loci differing between Tipple and Chevallier *Pseudomonas* isolates. Finally, cultivar-driven selection is important for plant health, with both cultivars showing a distinct preference for microbes selected by their genetic siblings in rhizosphere transplantation assays.

## Introduction

Plants grow in close association with diverse bacteria, fungi, protozoa, archaea and viruses that can influence the plant host in different ways by improving growth, protecting against pathogens or conferring adaptive advantages [1–3]. Every plant species possesses a unique microbial signature, or microbiome, which extends their capacity to adapt to the surrounding environment and enhances their ability to withstand biotic and abiotic stresses [4–6]. Distinct microbial communities inhabit different niches within the plant such as the phylosphere, endosphere and rhizosphere, with each of these micro-habitats providing a specific niche to which microbes must adapt and the composition of each compartment’s microbiome differing within an individual plant [7].

The rhizosphere and roots are hotspots for microbial life because of the availability of plant-derived root exudates. Plants secrete an average of 21% of their photosynthates into the rhizosphere, which triggers a specialised shift in microbial composition and increases microbial density 10 to 1000 times within the root influenced area [4, 8–10]. The microbiome of the rhizosphere constitutes the most populous and diverse set of microorganisms directly interacting with the plant. Consequently, the impact of this community on plant health, disease and productivity is highly significant [11].

Given that in general, soil nutrient content is quite low, access to plant-derived nutrients exacerbates the competition between microbial species for niche colonisation [12, 13]. In response, bacteria have developed numerous ecological traits to improve their rhizosphere colonisation competitiveness [14]. Among these, metabolic versatility is particularly important in *Pseudomonas*, a ubiquitous and important bacterial genus that lives in a wide range of environments [15, 16]. Plant-associated pseudomonads, such as the plant-growth promoting rhizobacterium (PGPR) *Pseudomonas fluorescens* non-specifically colonise a vast range of different plants and exert important impacts on plant health by enhancing nutrient availability, suppressing pathogens and priming the plant immune system [17]. *P. fluorescens* colonise diverse ecological niches and have correspondingly complex genomes, typically encoding ∼6000 genes with a remarkably high level of intraspecies diversity [18, 19]. Much of the *P. fluorescens* accessory genome is devoted to signal transduction, environmental interactions and specialised metabolism [17, 18].

Researchers have recently begun to uncover the molecular mechanisms underpinning the relationship between plants and their rhizosphere microbiomes. This includes key concepts such as the importance of starting soil microbiome [20], founder effects [20, 21] and local plant habitat [22, 23] on community composition, and the importance of microbiome composition for plant health and pathogen biocontrol [6, 24, 25]. Furthermore, rhizosphere community assembly is actively shaped by plant genetic determinants. For example, transcriptional rewiring during *Medicago* nodulation is influenced by plant genotype, which impacts the outcome of *Sinorhizobium* symbiosis [26]. Bulgarelli and co-workers showed that the barley genotype influences both root and rhizosphere microbiota, with variation seen for many microbes from diverse phyla [27]. Differential recruitment of taxonomically distinct rhizosphere bacteria was subsequently linked to a small number of loci in barley, with an NLR-like gene one of the main drivers of this phenomenon [28]. Another recent publication showed how a previously characterised receptor kinase, FERONIA [29] negatively modulates *P. fluorescens* colonisation of *Arabidopsis thaliana* by controlling the basal levels of reactive oxygen species [30].

Root exudate composition has also been shown to exert a significant effect on rhizosphere microbial recruitment. In *Avena barbata* bacterial soil isolates are differentially affected by plant growth, producing a split in the bacterial community based on their response to root growth over time [31]. Meanwhile, the secretion of coumarins [32] and root-derived triterpenes [33] in *A. thaliana* have been shown to play important roles in the rhizosphere microbiome assembly process, with the latter exerting strong impacts on the abundance and composition of Bacteroidetes and δ-proteobacteria [33]. While our understanding of rhizosphere microbiome assembly is rapidly advancing, at this stage we still know little about the impact of plant genotype on selection within individual soil species. Given the striking differences in plant-beneficial phenotypes seen for different members of the same PGPR species complex [19, 24], a better understanding of the principles underlying microbial genotype selection in the rhizosphere is a priority.

To address the effect that plant genotype exerts on microbial recruitment we analysed the rhizosphere microbiomes of two distinct barley cultivars with markedly distinct histories and genetic backgrounds. Chevallier is an English landrace cultivar first selected in the 1820s before breeding programs were established, whereas NFC Tipple (hereafter ‘Tipple’) is a modern cultivar released in 2004 by Syngenta Seeds, Ltd. [34]. Amplicon metabarcoding revealed distinct differences in the abundance of several rhizosphere and root-associated bacterial and fungal species between the two varieties, with Tipple recruiting significantly more *Pseudomonas* bacteria than Chevallier. This selection also manifested within the recruited *Pseudomonas* population, with distinct genotypic clustering by cultivar, leading to marked differences in the phenotypes of recruited isolates.

Cultivar-driven selection in barley appears to be primarily driven by root exudate composition. In general, the Tipple-associated *Pseudomonas* population was better adapted to grow on hexose sugars than the Chevallier population, aligning with a greater abundance of these molecules secreted from Tipple roots than Chevallier. Cultivar-driven selection was also observed for individual loci; both at a population level, where bacterial genes important for Tipple colonisation were preferentially selected by Tipple and likewise for Chevallier, and in differential gene transcription in the model organism *P. fluorescens* SBW25. Chevallier seedlings derived a growth advantage from the presence of microbes selected by either cultivar, with a slight preference for their ‘own’ microbiota. Conversely, while Tipple seedlings grew slightly better when colonised by Tipple-selected microbes, they responded poorly to the Chevallier-selected microbiota, supporting a link between cultivar-driven rhizosphere selection and plant health.

## Results

### Barley cultivar impacts root associated microbial communities

To gain an overview of the microbiome structure of the Chevallier and Tipple root systems, we used targeted amplicon sequencing of the 16S rRNA gene and ITS regions from gDNA samples taken from four-week-old barley plants (Figure 1, supplementary Figure 1). The relative abundance of amplicon sequence variants (ASVs) [35] for the 30 most represented bacterial genera are shown in Figure 1a. Overall, 63.3 % of all ASVs were Proteobacteria, 26.7 % Actinobacteria, 6.7 % Acidobacteria and 3.3 % Bacteroidetes.

**Figure 1.**
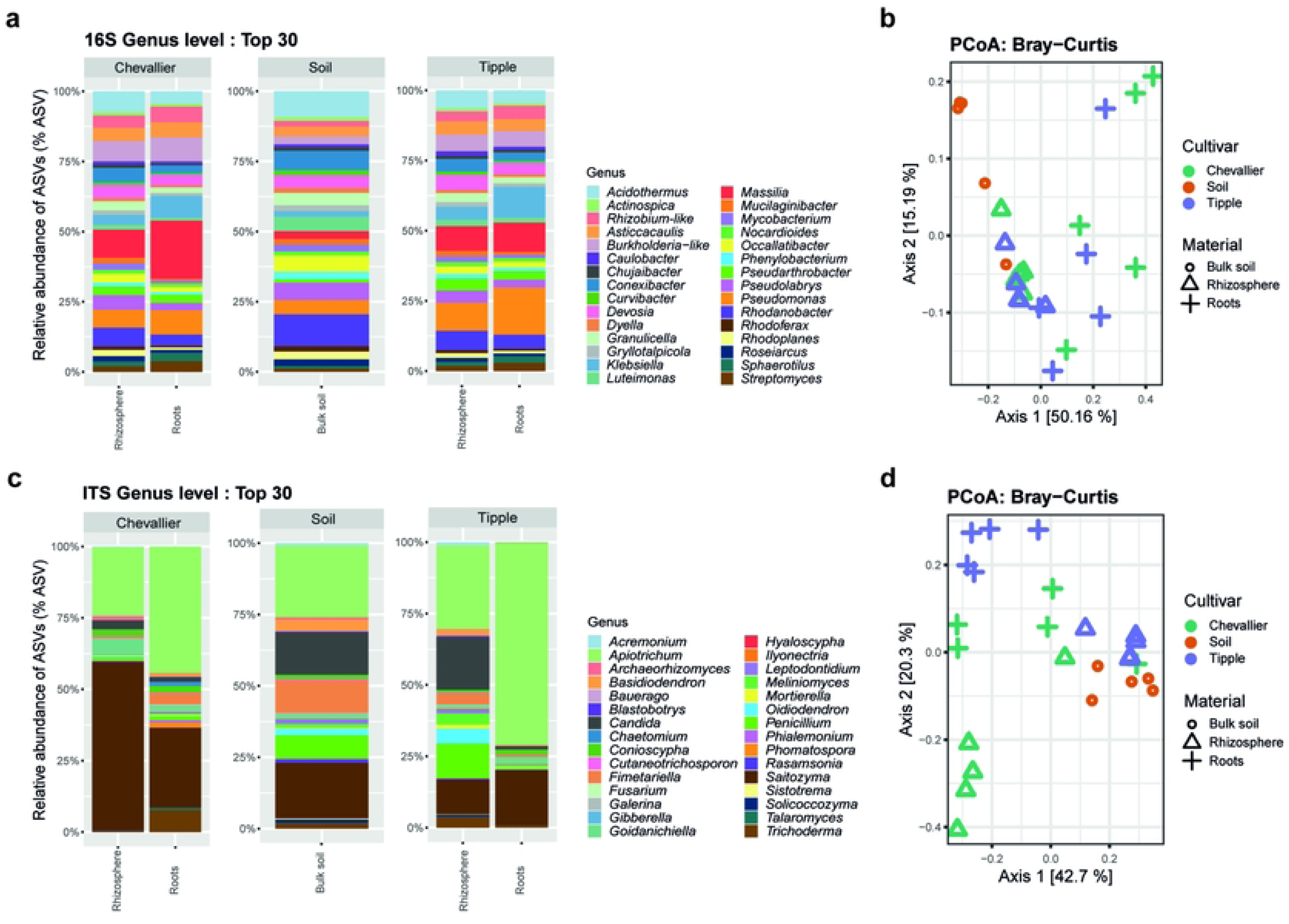
Metabarcoding analysis of the microbial communities associated with Chevallier, Tipple and bulk soil. **A.** Top 30 bacterial ASV community composition. **B.** PCoA showing beta-diversity with Bray-Curtis distance for bacterial community composition. **C.** Top 30 fungal ASV community composition. **D.** PCoA showing beta-diversity with Bray-Curtis distance for fungal community composition. Two plant compartments; root endosphere and rhizosphere, and the bulk soil were analysed. Five replicates were used per condition. In PCoA, green represents Chevallier, orange, bulk soil and purple, Tipple. Black hollow symbols represent plant compartment, round is bulk soil; triangle is rhizosphere and cross represents roots or endosphere.

In the bulk soil, the most abundant genera were *Rhodanobacter* (8.35 %), *Acidothermus* (6.82 %) and *Conexibacter* (5.11 %). The rhizosphere of the two cultivars (adonis test, *p*-value 0.001) and the different plant compartments (ANOSIM test, *p*-value 0.001) differed significantly in their bacterial composition. In the Tipple rhizosphere, *Pseudomonas* (7.55 %) and *Massilia* (6.62 %) presented the greatest relative abundance, while in the Chevallier rhizosphere the most abundant genera were *Massilia* (7.67 %) and *Acidothermus* (5.65 %), with *Pseudomonas* representing 4.87 % of total ASVs. In the Tipple root compartment, *Pseudomonas* (13.43 %), *Klebsiella* (8.82 %) and *Massilia* (8.36 %) were enriched, while for Chevallier the most abundant genera were *Massilia* (17.03 %), *Pseudomonas* (7.15 %), members of the Burkholderiaceae family (7.04 %) and *Klebsiella* (6.49 %).

Next, Principal Coordinates Analysis (PCoA) using Bray-Curtis distance measurement was used as a measure of the variation of taxonomic profiles between samples (Figure 1b). The first component accounted for about 50 % of the variance, with samples clustering both by compartment (rhizosphere, root, or bulk soil) and host cultivar. Bacterial composition between cultivars was significantly different (adonis test, *p-*value 0.001) as well as between roots, rhizosphere and bulk soil (ANOSIM test, *p-*value 0.001). Within individual cultivars, both the Chevallier (ANOSIM test, *p-*value 0.01) and Tipple (ANOSIM test, *p-*value 0.01) root and rhizosphere communities were statistically distinct from one another. Finally, the Chevallier and Tipple rhizosphere communities were also significantly different (adonis test, *p-*value 0.01).

*Saitozyma* and *Apiotrichum* were the most abundant plant-associated fungal genera, together accounting for up to 90 % of the ASV abundance in some samples (Figure 1c). These two genera were also highly abundant in bulk soil, with *Candida* (14.78 %), *Fimetariella* (11.40 %) and *Penicillium* (8.17 %) the next most prevalent genera. The Chevallier rhizosphere community differed markedly from bulk soil, with a very high abundance of *Saitozyma* and an increase in *Goidanichiella* (5.78 %). By contrast, the Tipple rhizosphere displayed a very similar profile to bulk soil, with *Candida* (18.39 %) and *Penicillium* (12.01 %) among the most abundant genera. The Tipple root compartment displayed a dramatic reduction in fungal diversity, with most samples overtaken by *Apiotrichum* and to a lesser extent *Saitozyma.* This reduction in diversity was less marked for the Chevalier root system. Here, *Trichoderma* (7.66 %) was the third most abundant genus in the root compartment. Bray-Curtis beta-diversity showed clear clustering of samples both by host cultivar and by compartment (Figure 1d), with significant differences observed between both host cultivars (ANOSIM test, p-value 0.005) and compartments (adonis test, p-value 0.002).

To assess the species richness across barley cultivars and plant compartments, we calculated alpha-diversity for bacterial and fungal communities (Supplementary Figures 1 and 2). For the bacterial population, alpha diversity decreased progressively from bulk soil to rhizosphere to the root compartment. As shown by Shannon index, significant differences were observed between the rhizospheres of both cultivars and the bulk soil, but not between the two rhizosphere communities (Supplementary Figure 1a). Differences in the observed species richness (ANOVA test, p-value 0.045) were seen for the root endosphere communities of the two cultivars (Supplementary Figure 1b), but not for Shannon index. For Chevallier, both observed richness (Krustal test, p-value 0.03) and Shannon index (Krustal test, p-value 0.009) were significantly different between plant compartments (Supplementary Figure 1c), while Tipple presented a marked difference between rhizosphere and root bacterial communities by Shannon index (Krustal test, p-value 0.009) but not observed richness (Supplementary Figure 1d).

Alpha diversities of the fungal population were significantly different between the Chevallier rhizosphere and soil for both observed richness (TukeyHSD test, p-value 9.6E^-5^) and Shannon index (Pairwise Wilcox test, p-value 0.024) and between Chevallier and Tipple rhizospheres (observed, TukeyHSD test p-value 0.001; Shannon, Pairwise Wilcox test p-value 0.024). No other significant differences were observed, (Supplementary Figure 2b, 2c), apart from a highly significant change in alpha-diversity between the Tipple root and rhizosphere populations (observed, ANOVA test p-value 0.004; Shannon, ANOVA test p-value 0.0002) (Supplementary Figure 2d).

These experiments suggest that (i) ecological niche plays a fundamental role in microbial community assembly and structure; (ii) species richness progressively decreases from the soil to the root endosphere; (iii) barley genotype affects the assembly of plant-interacting microbial consortia; (iv) Chevallier substantially modifies the fungal composition of its rhizosphere, while Tipple does not, and (v) key bacterial genera are preferentially enriched by Tipple (e.g. *Pseudomonas*) or Chevallier (e.g. *Massilia*) root environments.

### Rhizosphere Pseudomonas genotypes cluster according to plant genotype

To interrogate the cultivar-specific differences more closely, we next examined the *Pseudomonas* populations associated with Chevallier and Tipple. While several fungal and bacterial genera displayed larger cultivar-specific differences, *Pseudomonas* spp. have important effects on plant growth and health, are easy to isolate and are genetically tractable, making them a good model for downstream molecular analysis. We used CFC selective agar to randomly isolate 239 bacterial strains from barley roots, alongside 31 bulk soil isolates. Genotyping based on the *gyrB* housekeeping gene [36, 37] identified most isolates as *Pseudomonas* spp., predominantly from the *P. fluorescens* complex (Figure 2). 21 isolates were determined to be species other than *Pseudomonas* and were excluded from further analysis. The resulting phylogenetic tree showed clear patterns of clustering, corresponding both to the experiment (and hence soil batch) but also to barley cultivar. The bulk soil isolates (representing a subset of the initial *Pseudomonas* population before barley planting), were distributed evenly throughout the tree. We then assessed the overall diversity present in the *Pseudomonas* population by computing the estimated divergence in *gyrB* sequences using the Tamura-Nei model [38]. By this calculation, the bulk soil isolates showed the greatest sequence divergence (0.1509), followed by the Chevallier isolates (0.1423), with the Tipple population showing the least sequence divergence (0.1225). The total population had a distance index of 0.1371.

**Figure 2.**
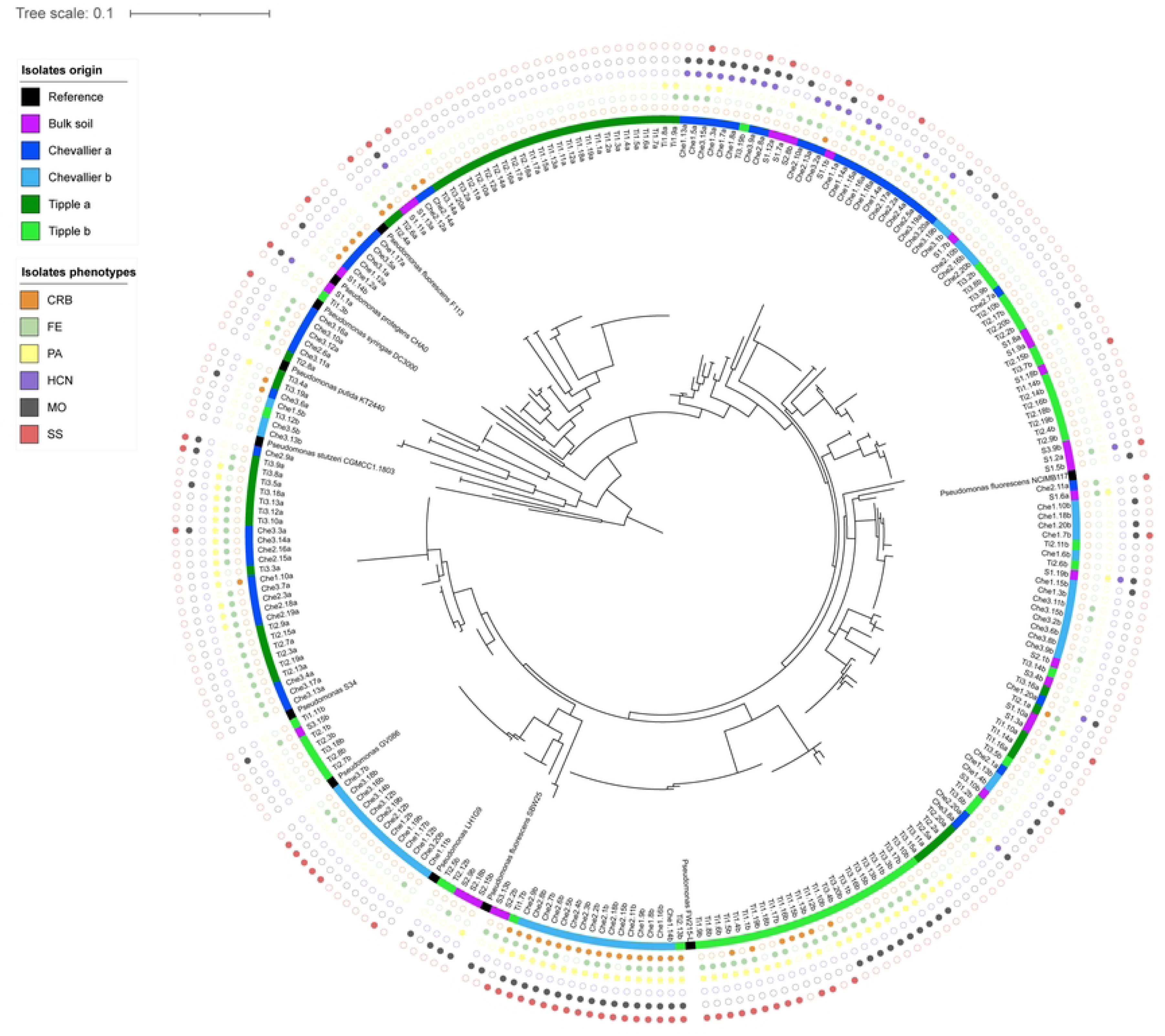
*gyrB* phylogeny of the Chevallier and Tipple rhizosphere *Pseudomonas* isolates. Two independent datasets, denoted with suffixes a and b, are included in the analysis. The phylogenetic tree is based on 260 partial sequences of the *gyrB* gene from sequenced and type strains and was constructed using the ML method and Tamura-Nei model with 1000 bootstrap value. The isolation origin of samples is indicated by the coloured ring closest to the tree. Reference strains are based on publicly available genome sequences. Filled/empty circles in the outer rings denote the presence/absence respectively of the ecological traits shown in the ‘Isolates phenotypes’ key.

We next examined the extent to which the apparent genotypic selection within the rhizosphere *Pseudomonas* community translated into differences in the distribution of phenotypic traits. Isolates were tested for Congo Red binding (CRB) -used here as a proxy for their biofilm forming capacity-, fluorescence emission (FE) -to measure siderophore production-, protease activity (PA), hydrogen cyanide (HCN) production, swarming motility (MO) and suppression of the Actinomycete *Streptomyces venezuelae* (SS). Ordinal values were then assigned to each trait (Figure 2). The Chevallier isolates contained a higher percentage of isolates with high CRB, MO or HCN scores, whereas Tipple isolates scored lower for almost all the tested traits (Supplementary Figure 3). A striking degree of clustering was observed between phenotypic traits and the distribution of *Pseudomonas* genotypes in our phylogenetic tree (Figure 2).

### Chevallier and Tipple differ in root exudates composition

We hypothesised that the differences observed in microbiome composition and microbial genotype distribution between Chevallier and Tipple could be driven by differences in the secretion of root exudates. To test this, we used luminescent biosensors [39, 40] to examine the secretion of specific groups of metabolites from the roots of barley seedlings. Fructose, C4-dicarboxylates, sucrose and phenylalanine biosensors were selected to represent key groups of molecules often found in the rhizosphere; sugars, organic acids and amino acids [41], and plants were evaluated at 2-, 5- and 7- days post-inoculation (dpi). Significant differences between cultivars were seen for the fructose biosensor, with especially high expression in Tipple at 2 dpi (Figure 3a).

**Figure 3.**
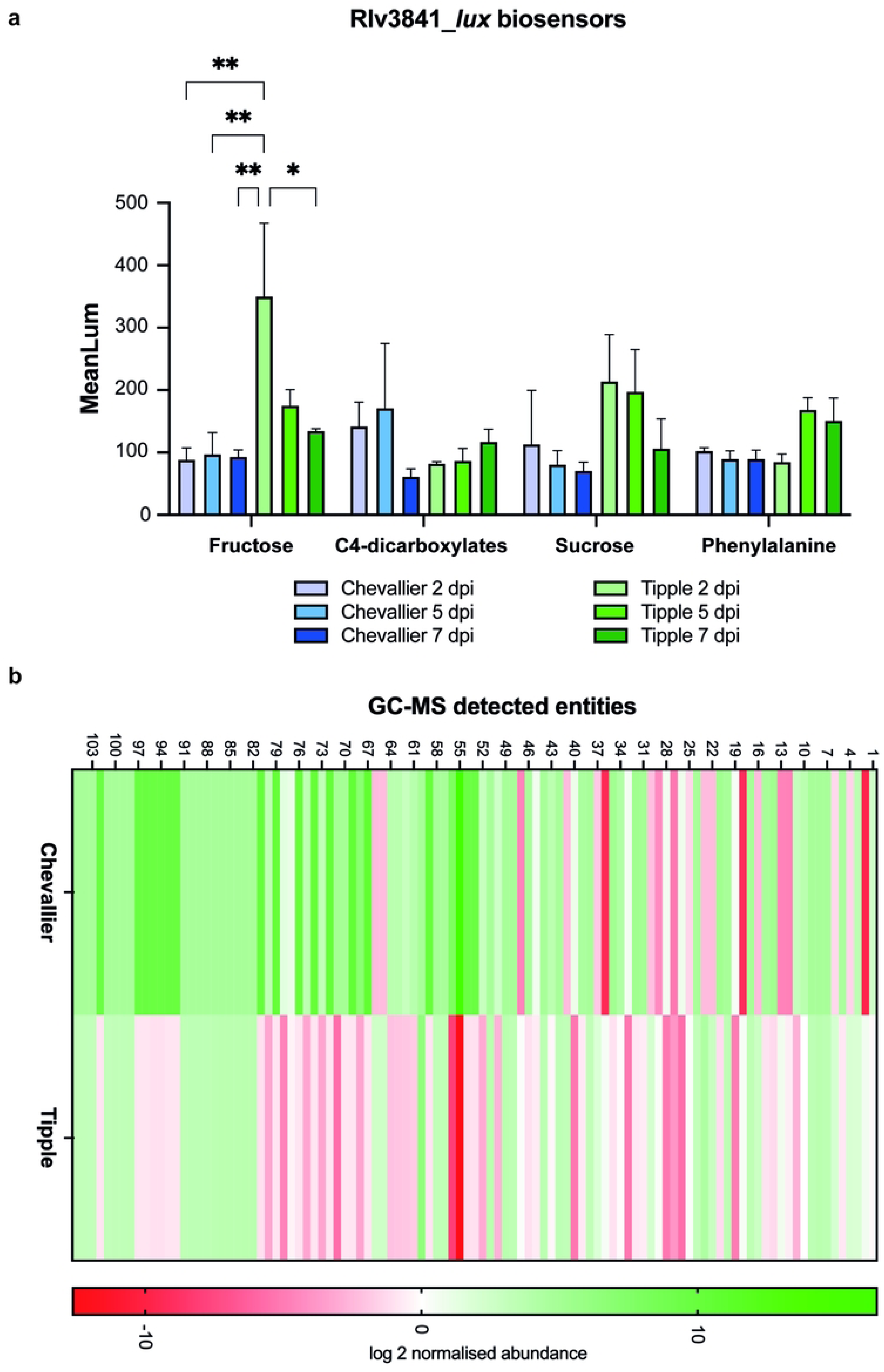
Root exudate analysis of Chevallier and Tipple. **A.** *In planta* temporal screening of root metabolites produced by Chevallier and Tipple seedlings with luminescent biosensors. Fructose, C4-dicarboxylates, sucrose and phenylalanine biosensors based on WT Rlv3841 [39] were inoculated in the roots of 2 days old Chevallier and Tipple seedlings. Images were acquired at 2, 5 and 7 dpi and 3 plants were used per condition. Mean values of luminescence intensity are shown in the graph. Error bars represent Standard Error of the Mean, p-values were calculated by Tukey’s multiple comparison test and asterisks indicate p < 0.05 (*), 0.01 (**). A representative graph is shown from three independent experiments. **B.** Chevallier and Tipple root exudates metabolites detected by GC-MS (gas chromatography mass spectrometry). The heat map shows the overall composition of root exudates identified metabolites presenting a log2-fold change ≥ 2 in at least one of the cultivars studied.

We then explored the root exudates diversity directly using gas chromatography–mass spectrometry (GC-MS). Chevallier and Tipple barley plants were grown for 3 weeks under axenic conditions and their root exudates were extracted and analysed. A total of 105 entities with a log_2_-fold change > 2 were detected by GC/MS (Figure 3b, Supplementary Table 1). Compounds 1-59 showed a match in at least one of the compound libraries used, whereas 60 to 105 could not be identified. Overall, a greater abundance of the unidentified metabolites was detected in the Chevallier exudates compared to Tipple. Two striking differences were seen between the two cultivars. First, a derivative of phosphoric acid was massively abundant in Chevallier exudates (log_2_-fold change 16.474) and absent in Tipple (log_2_-fold change −12.613). Second, _D_-glucose, whose log_2_ abundance was 4.389 in Tipple, was much less abundant (−2.405) in Chevallier exudates.

Overall, these data suggest that the Tipple rhizosphere is enriched in C_6_/C_12_ sugars, such as fructose and glucose. In contrast, Chevallier exudates contain a more diverse exudate composition and a greater abundance of unidentified molecules.

### Primary carbon metabolism is central to Pseudomonas spp. cultivar adaptation

To examine the link between the exudate profiles of Chevallier and Tipple and the rhizosphere *Pseudomonas* population, we tested the ability of the *Pseudomonas* isolate library to grow on different carbon sources. Comparing the final OD_600_ values for 24 hour-grown cultures, we observed two intriguing trends (Figure 4). When grown on acetate as the sole carbon source (Figure 4a), several Chevallier (7/60), but no Tipple isolates attained a final OD_600_ >0.3. Conversely, when grown on glucose (Figure 4b), 8 Tipple isolates reached a final OD_600_ >0.25, but no Chevallier isolates, supporting the hypothesis that root exudate differences may underpin the observed selection of *Pseudomonas* genotypes (Figure 2).

**Figure 4.**
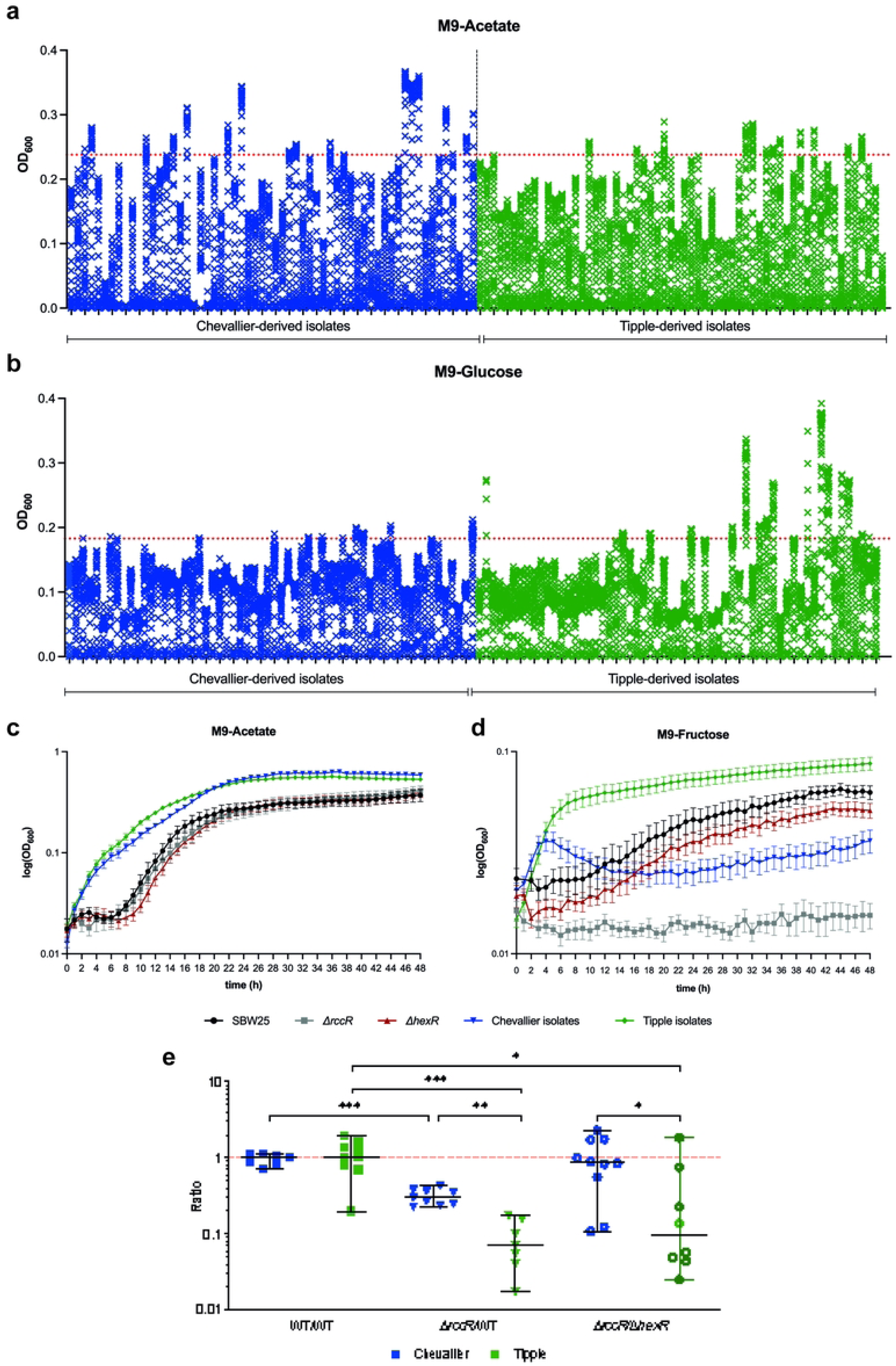
Carbon regulation is a key determinant for rhizosphere microbe selection. **A., B.** Rhizosphere *Pseudomonas* isolate growth over 48 h in M9 medium supplemented with 0.4 % Acetate (**A**) or 0.4 % Glucose (**B**). Each cross in a vertical stack denotes the OD_600_ value at sequential time points from 0 to 48 h. Values were normalised to the lowest value of each dataset. Chevallier isolates are coloured blue and Tipple isolates green. The red line represents the 75th percentile of all 48 h OD_600_ values. **C., D.** Growth curves for the 42 rhizosphere *Pseudomonas* isolates selected for whole genome sequencing over 48 h in M9 medium supplemented with 0.4 % acetate (**C**) or 0.4 % fructose (**D**). SBW25 WT is shown in black, SBW25 Δ*rccR* in grey, SBW25 Δ*hexR* in red, pooled Chevallier isolate values in blue and pooled Tipple isolates in green. Values for Chevallier and Tipple isolates are represented as the mean values of all the isolates tested. Std. errors shown in each case. **E.** Chevallier/Tipple rhizosphere colonisation competition assays. The graph shows the ratios of SBW25 WT and Δ*rccR* to SBW25-*lacZ* or SBW25 Δ*rccR*-*lacZ* to Δ*hexR*. CFUs recovered from the rhizospheres of Chevallier (blue) and Tipple (green) barley cultivars at 5 dpi. Each dot represents the ratio of CFUs recovered from an individual plant. 8-10 plants were used per condition and p-values were calculated by Mann-Whitney U test, asterisks indicate p < 0.05 (*), 0.01 (**) or 0.001(***). Experiments were repeated at least twice, and representative graphs are shown here.

To investigate further, a subset of 42 isolates was selected for more detailed analysis. 19 strains were chosen based on a combination of growth characteristics (Figure 4a-b) and phylogenetic distribution to ensure maximum genetic diversity (Figure 2). A further 23 isolates were then selected at random to reduce bias within the sample set. These isolates were again interrogated for growth in defined media. Alongside acetate, we selected fructose as a hexose sugar abundant in many root exudates [42, 43], including Tipple. No obvious differences in growth rates were observed between Tipple or Chevallier isolates when grown on acetate (Figure 4c). However, when the isolates were grown on fructose (Figure 4d) a clear difference emerged between the Tipple and Chevallier isolates, with the Tipple-derived community growing faster and to a higher cell density than strains isolated from Chevallier.

Finally, to study the effect of differential root exudation on microbial colonisation, we performed rhizosphere competition assays using Tipple and Chevallier, and two well-characterised metabolic mutants of *P. fluorescens* SBW25: Δ*rccR* and Δ*hexR* [44]. Δ*rccR* displays a pronounced growth defect on compounds that enter the Krebs Cycle via the Entner Doudoroff (ED) pathway. Conversely, Δ*hexR* struggles to grow on Krebs Cycle intermediates and two-carbon molecules such as acetate ([44], Figure 4d). While the Δ*rccR* mutant was less competitive than WT in both cultivars, its competitive disadvantage was significantly greater in the Tipple rhizosphere. Δ*rccR* and Δ*hexR* showed similar fitness in the Chevallier rhizosphere, but Δ*rccR* was strongly outcompeted by Δ*hexR* in Tipple (Figure 4e). Together, these data support the hypothesis that barley cultivars select *Pseudomonas* genotypes that can most effectively metabolise their root exudates.

### Different barley rhizospheres influence the abundance of specific bacterial loci

Next, we examined the effect of cultivar-specific selection on the genomes of the rhizosphere Pseudomonas community. Our 42 selected isolates were Illumina whole genome sequenced and reciprocal BLAST-searched for the presence of loci potentially involved in plant colonisation, using the model organism SBW25 as a reference genome. In the first instance, this included 410 previously described plant-induced *Pseudomonas* genes, encoding transporters, biofilm regulators, chemotaxis proteins or siderophore synthases [45] alongside substrate transporters and transcriptional regulators - two important traits relevant to niche adaptation [13, 44] (see materials and methods). Using a Chi-Square analysis, we identified 51 genes whose distributions significantly differed in the Chevallier and Tipple genomes (Figure 5a). Some genes were abundant in Tipple isolates but scarce in Chevallier strains, e.g., the sugar ABC transporter gene *PFLU_2583*. Likewise, others were found at higher rates in Chevallier isolates, such as *PFLU_0315*, encoding a GABA transporter. The largest group of differently abundant loci were found in the KEGG categories of signalling and cellular processes, including metabolism and signal transduction (23 % of selected genes), and transporters (20 %) (Figure 5b).

**Figure 5.**
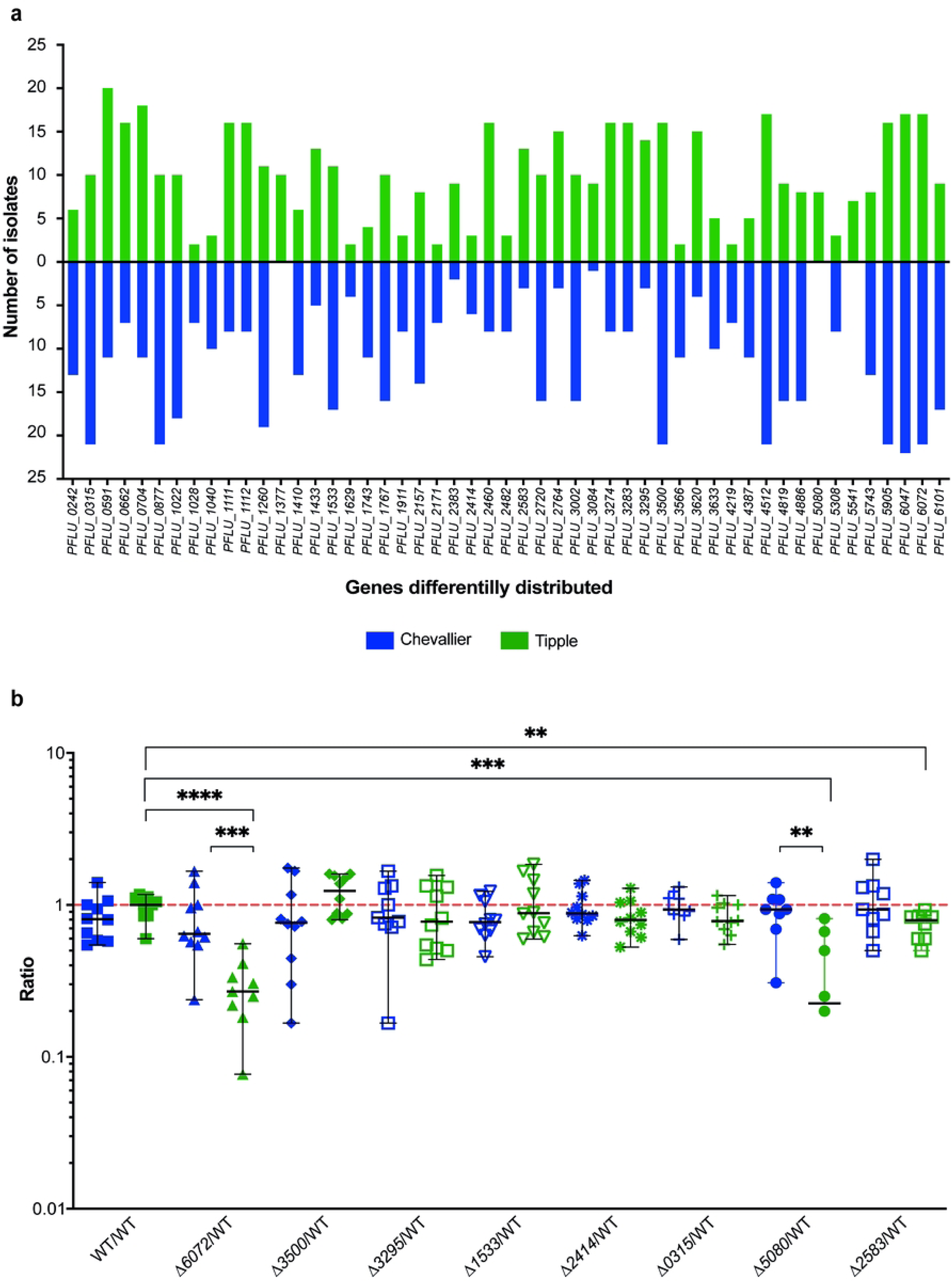
Cultivar-driven gene selection within the barley rhizosphere *Pseudomonas* population. **A.** Distribution of genes differentially present in the sequenced *Pseudomonas* isolates from Chevallier and Tipple. The bars show the number of strains within the sequenced population where reciprocal BLAST suggests a particular gene is present. The SBW25 gene designation for each locus is given on the X-axis. Chevallier isolates are shown in blue and Tipple, in green. All the genes represented here were significantly differently abundant (p < 0.05) between the two cultivars according to a Chi-Square test. **B.** Rhizosphere colonisation competition assay. The graph shows the ratios of differentially distributed mutants to SBW25-*lacZ* CFUs recovered from the rhizospheres of Chevallier (blue) and Tipple (green) barley cultivars. Each dot represents the ratio of CFUs recovered from an individual plant. p-values were calculated by Mann-Whitney U test and asterisks indicate p < 0.05 (*), 0.01 (**), 0.001(***) or 0.0001(****). The experiment was repeated three times and a representative graph is shown here.

To investigate the biological roles of the potential cultivar-selected genes, we produced non-polar deletions of eight genes in SBW25. Genes were selected based on their predicted biological role and the extent that their frequency differed between the two cultivars. The deleted genes and their predicted functions are listed in Table 1. Growth assays in complex (KB and LB) and defined minimal media showed that Δ*6072* exhibited compromised growth in pyruvate, glucose, or glycerol minimal media (Supplementary Figure 4) implicating this predicted LysR-family regulator gene in the control of primary carbon metabolism.

**Table 1.** Cultivar-selected genes deleted in SBW25

| Gene ID | Predicted function of encoded protein | More abundant in: |
| --- | --- | --- |
| *PFLU_0315* | GABA permease: *gabP* | Chevallier |
| *PFLU_1533* | LysR-family transcriptional regulator | Chevallier |
| *PFLU_2414* | Iron siderophore sensor | Chevallier |
| *PFLU_6072* | LysR-family transcriptional regulator | Chevallier |
| *PFLU_2583* | Putative rhizopine-binding protein | Tipple |
| *PFLU_3295* | GntR family transcriptional regulator: *vanR* | Tipple |
| *PFLU_3500* | C4-dicarboxylate transporter | Tipple |
| *PFLU_5080* | Hypothetical protein | Tipple |

Chevallier and Tipple rhizosphere competition assays were then carried out for the eight mutants (Figure 5b). Δ*6072* showed an impaired colonisation phenotype versus WT SBW25 in both cultivars, which was highly significant in the Tipple rhizosphere. Given the growth impairment of this mutant on glucose, this result aligns with the observation that the Tipple rhizosphere is enriched in glucose and similar molecules. Δ*2583* also exhibited a fitness penalty, but only on Tipple plants. As mentioned above, *PFLU_2583* encodes a putative sugar transporter and is found in greater abundance in Tipple rather than Chevallier-associated isolates. Finally, Δ*5080* also displayed a compromised colonisation ability in Tipple rhizospheres only. These results support the idea that specific genetic traits are selected by root exudates in a cultivar-specific manner, and in turn support rhizosphere colonisation on those specific cultivars.

### Cultivar specific genes were differentially expressed in SBW25

Next, we investigated to what extent this genetic selection overlaps with differential gene expression in the barley rhizosphere. To address this, SBW25 mRNA abundance was studied by RNA-seq in the Tipple and Chevallier rhizospheres, 1 and 5 dpi (Figure 6a-f). A total of 2,158 genes were differentially expressed (log_2_-fold ≤ −1 or ≥1 and *p*-value ≤ 0.05) compared to the cell-culture control in the rhizosphere of Chevallier at 1 dpi (Figure 6a), increasing slightly to 2,515 at 5 dpi (Figure 6b). 2,039 genes were differentially expressed in the Tipple rhizosphere at 1 dpi (Figure 6c), of which 950 were upregulated and 1,089, downregulated. At 5 dpi this increased to 2,470 genes (Figure 6d).

**Figure 6.**
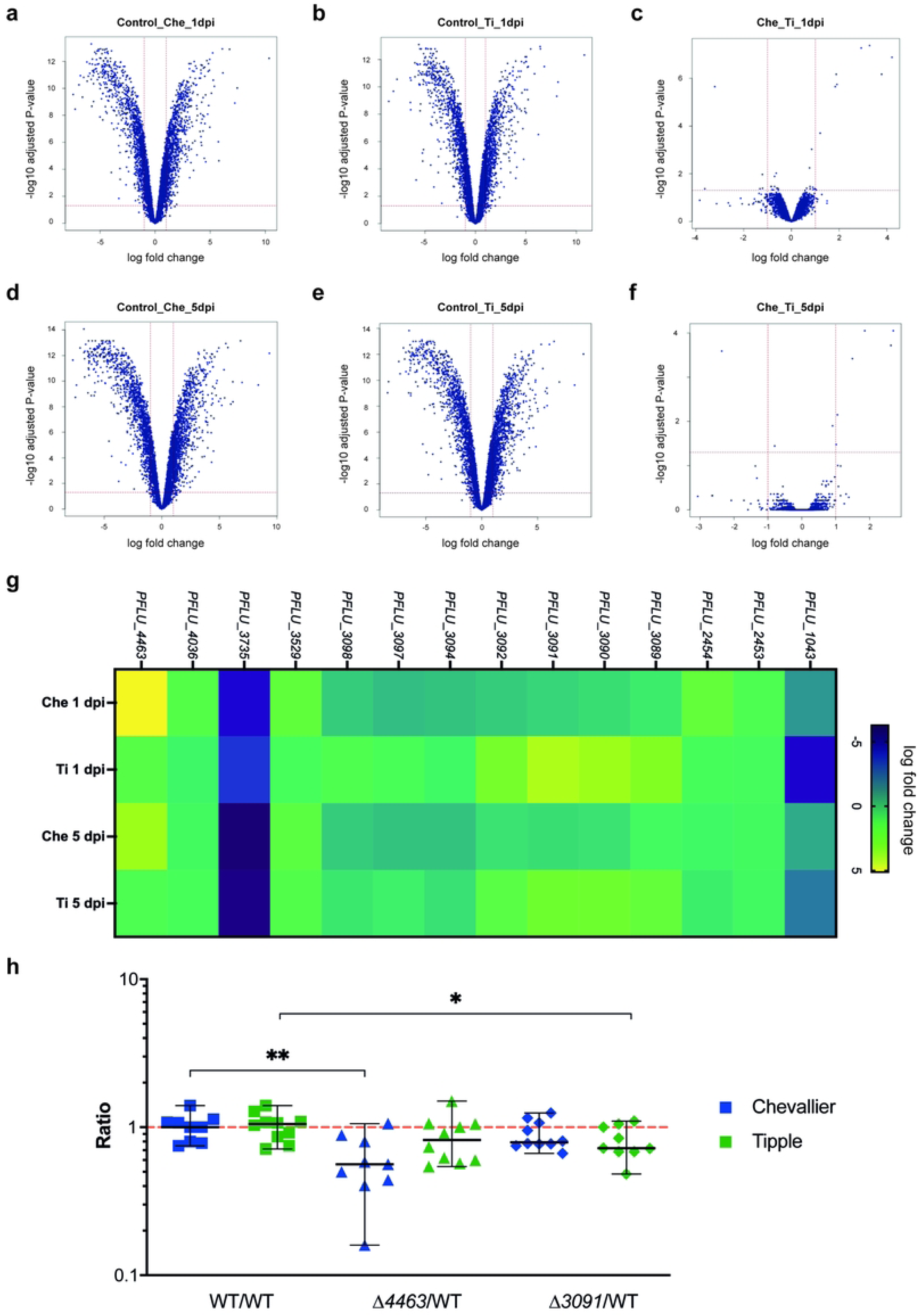
SBW25 gene expression is affected by plant cultivar. **A.** Volcano plot comparison of SBW25 transcript abundance between liquid culture control and Chevallier rhizosphere at 1 dpi. **B.** Volcano plot comparison of SBW25 transcript abundance between liquid culture control and Tipple rhizosphere at 1 dpi. **C.** Volcano plot represents the pairwise comparison of transcript abundance between SBW25 extracted from the Chevallier rhizosphere and from the Tipple rhizosphere at 1 dpi. **D-F.** Volcano plots comparing transcript abundance at 5 dpi for the samples shown in **A-C**, respectively. **G.** Heat map showing significantly differentially expressed genes in SBW25 between the Chevallier and Tipple rhizosphere samples at 1 dpi and 5 dpi (>1 log2- fold change from liquid culture control, p-values < 0.05). **H.** Rhizosphere colonisation competition assay. The graph shows the ratios of Δ*3091* and Δ*4463* to SBW25-*lacZ*. CFUs recovered from the rhizospheres of Chevallier (blue) and Tipple (green) barley cultivars. Each dot represents the ratio of CFUs recovered from an individual plant. 8-10 plants were used per condition and p-values were calculated by Mann-Whitney U test, asterisks indicate p < 0.05 (*) and 0.01 (**). Experiment was repeated twice, and a representative graph is shown here.

In contrast to the high number of rhizosphere-regulated genes, pair-wise comparisons between the Chevallier and Tipple rhizosphere samples (Figure 6e and f) showed a striking similarity in SBW25 gene expression. At 1 dpi, just 14 genes were differentially expressed between the two cultivars, with only 7 genes presenting a significant fold-change after 5 dpi (Figure 6g). Genes whose mRNA abundance significantly differed between the Tipple and Chevallier rhizospheres are listed in Table 2. Curiously, most Tipple upregulated genes were in the same gene cluster, despite forming at least four distinct operons (Supplementary Figure 5). No significant differences in gene expression were observed for the differently selected genes discussed above (Figure 5, Supplementary Table 2). However, half of these loci were shown to be upregulated in the rhizosphere relative to liquid culture, independent of cultivar (Supplementary Table 2).

**Table 2.**
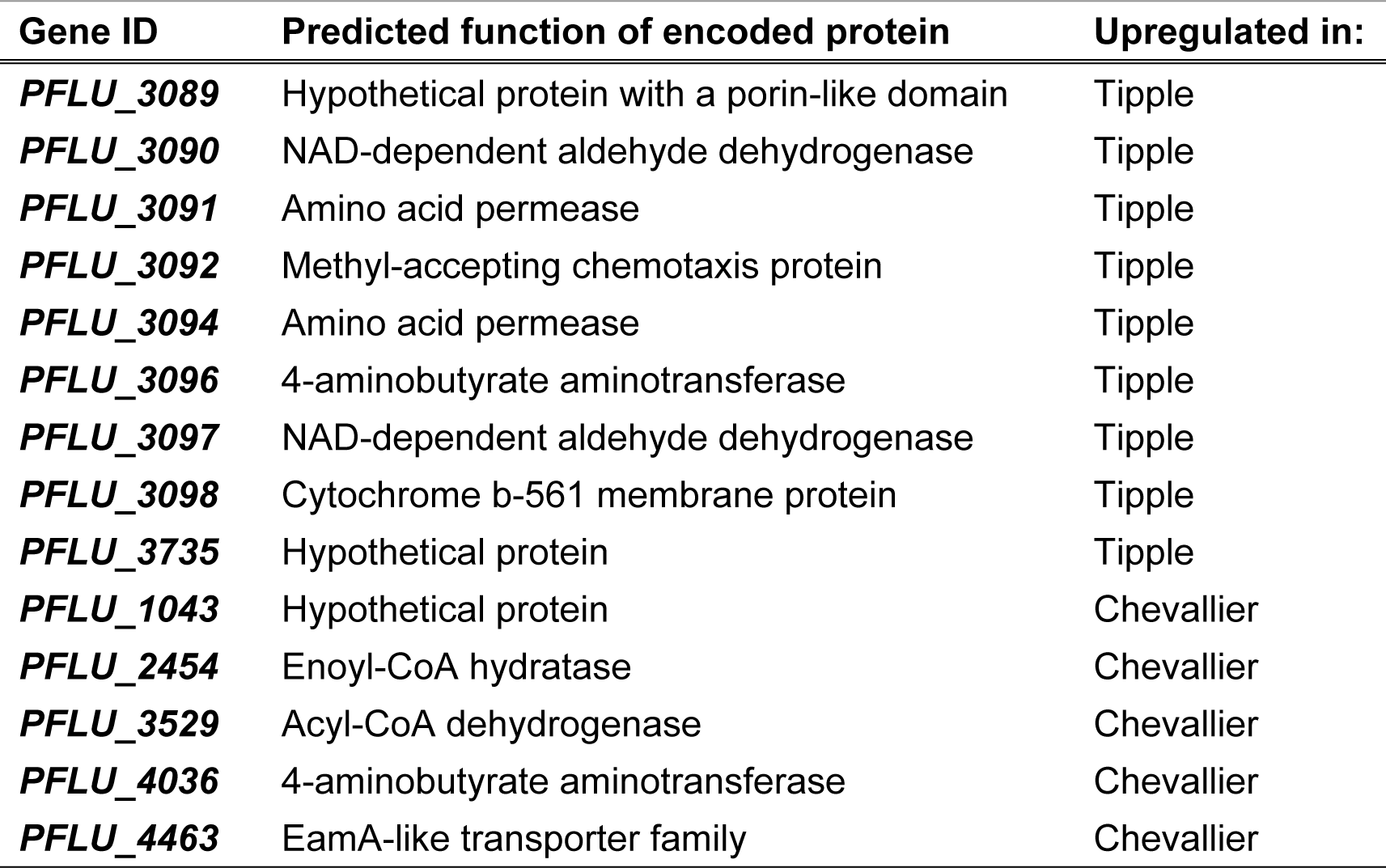
Predicted functions of genes differentially expressed in SBW25 between the Chevallier and Tipple rhizospheres. The chromosomal organisation of genes *PFLU_3089* to *PFLU_3098* is represented in Supplementary figure 3.

| Gene ID | Predicted function of encoded protein | Upregulated in: |
| --- | --- | --- |
| *PFLU_3089* | Hypothetical protein with a porin-like domain | Tipple |
| *PFLU_3090* | NAD-dependent aldehyde dehydrogenase | Tipple |
| *PFLU_3091* | Amino acid permease | Tipple |
| *PFLU_3092* | Methyl-accepting chemotaxis protein | Tipple |
| *PFLU_3094* | Amino acid permease | Tipple |
| *PFLU_3096* | 4-aminobutyrate aminotransferase | Tipple |
| *PFLU_3097* | NAD-dependent aldehyde dehydrogenase | Tipple |
| *PFLU_3098* | Cytochrome b-561 membrane protein | Tipple |
| *PFLU_3735* | Hypothetical protein | Tipple |
| *PFLU_1043* | Hypothetical protein | Chevallier |
| *PFLU_2454* | Enoyl-CoA hydratase | Chevallier |
| *PFLU_3529* | Acyl-CoA dehydrogenase | Chevallier |
| *PFLU_4036* | 4-aminobutyrate aminotransferase | Chevallier |
| *PFLU_4463* | EamA-like transporter family | Chevallier |

Two of the most differently regulated genes in the rhizospheres of the two cultivars, *PFLU_3091* and *PFLU_4463*, were selected for further characterisation. The predicted amino acid permease gene *PFLU_3091* is highly upregulated in the Tipple rhizosphere compared to Chevallier, while the EamA domain-containing gene *PFLU_4463* is downregulated in Tipple. Non-polar deletions of both genes were produced in SBW25, and the relative competitiveness of the two mutants assessed in the Tipple and Chevallier rhizospheres (Figure 6h). The Δ*4463* mutant was strongly compromised for Chevallier rhizosphere colonisation, while the Δ*3091* strain was less strongly, but still significantly compromised, in the Tipple rhizosphere. These results support roles for both *PFLU_3091* and *PFLU_4463* in cultivar-specific colonisation of the barley rhizosphere by *P. fluorescens*.

### Rhizosphere selected microbes induce cultivar and microbiome-specific plant growth

Finally, to examine the consequences of microbial selection for plant growth and health, we conducted a rhizosphere cross-inoculation experiment (Figure 7). Chevallier and Tipple seedlings were axenically inoculated with either a Chevallier rhizosphere microbiome extract (CRh) or a Tipple rhizosphere extract (TRh) and with synthetic communities (SynComs) of Chevallier- (CPs) or Tipple- derived *Pseudomonas* (TPs), then their dry weights were recorded after three weeks of growth. Tipple plants showed a small reduction in dry mass of −7 +/− 7.534 % when inoculated with CRh (TxCRh) in comparison to a Tipple uninoculated control, against a slight increase in mass (+12 +/ 12.610 %) when grown with their native rhizosphere community (TxTRh). Conversely, Chevallier showed strong positive responses to the rhizosphere extracts of both cultivars, with a mild preference for its own microbiome. Similarly, Chevallier inoculated with either CPs or TPs showed a growth increase (+15 +/− 12.14 % and +20 +/− 9.806% dry mass respectively), Tipple responded negatively to both SynComs. Together, these results suggest that Tipple is less capable than Chevallier of utilising a non-host rhizosphere microbiome. Furthermore, *Pseudomonas* SynComs promote Chevallier, but not Tipple plant growth.

**Figure 7.**
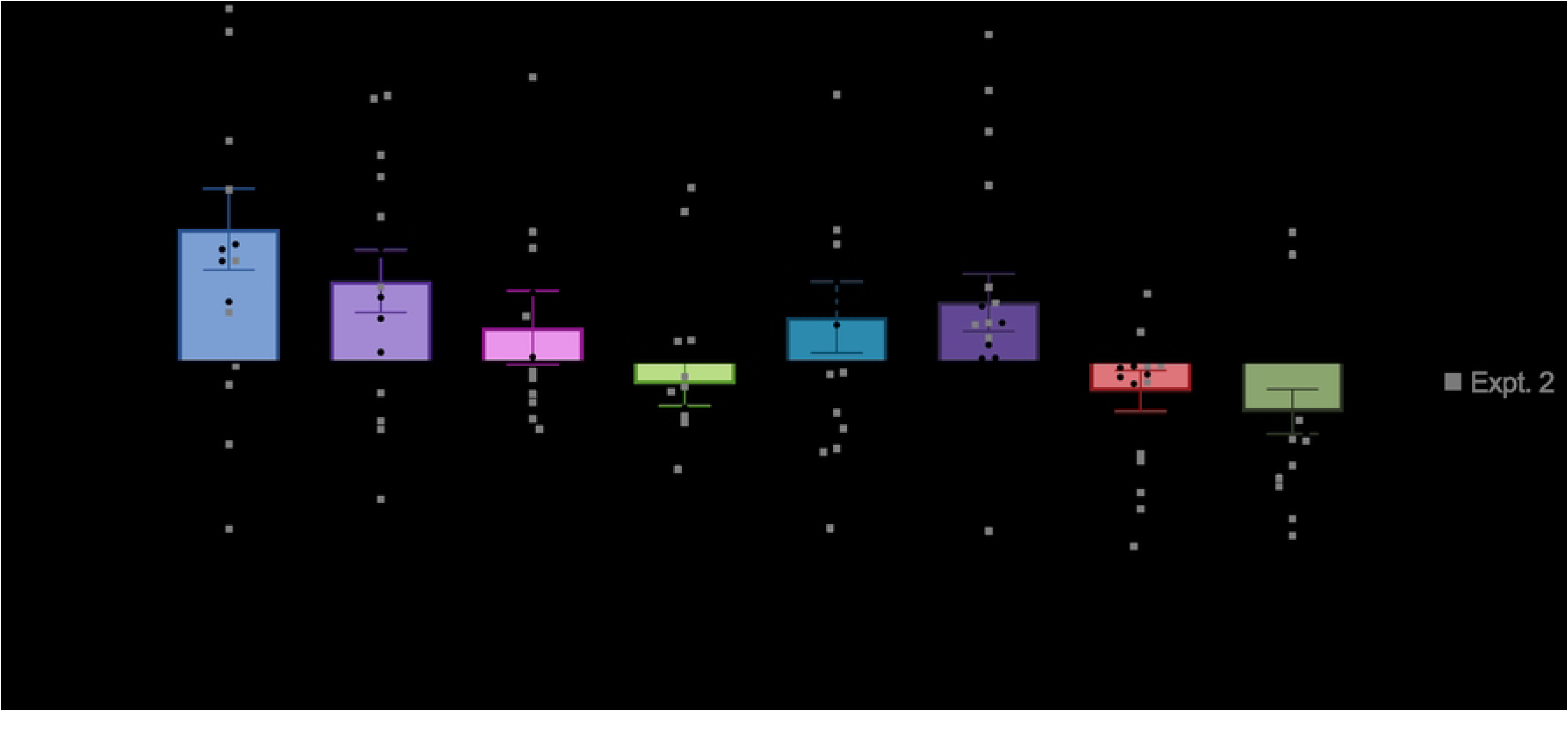
Cross-inoculation assay between Chevallier and Tipple. Chevallier (C) or Tipple (T) seedlings were inoculated under controlled conditions either with a rhizosphere extract (CRh/TRh) or with a SynCom of rhizosphere *Pseudomonas* (CPs/TPs). The graph summarises the dry weights (DW) of inoculated plants at 3 weeks post inoculation relative to the uninoculated controls. Two biologically independent repetitions of the experiment are represented side by side and up to 10 plants were used per condition. Data are presented as mean +/− std error. p-values were calculated by Tukey’s multiple comparison test and asterisks indicate p < 0.05 (*), 0.01(**).

## Discussion

Modern cereal varieties, such as Tipple, have been intensively bred for positive agricultural traits including high yield and seed starch content, short straw and good malting properties. However, while incorporation of these positive traits has substantially improved yield and crop quality, the wider effect of these changes on plant physiology and ecological impact are poorly understood. Consequently, there has been considerable recent effort to unravel the effect of plant genotype on the composition of the associated microbiota [6, 27, 30, 33, 46, 47].

While several studies have demonstrated the importance of plant genes [30] or metabolic pathways [33] for microbial recruitment, resolution within the microbiome itself has typically been restricted to genus (or occasionally species) level changes for whole microbiome studies. Studies using synthetic microbial communities [6, 46] enable interactions between microbes and hosts to be studied in greater detail, however these communities typically only contain a few representatives of each genus, limiting their utility to probe the importance of bacterial genotype in host colonisation. This issue also cannot be fully addressed by molecular analyses of bacterial colonisation pathways [44, 47], where the importance of individual loci in the highly complex plant environment is often difficult to extrapolate from laboratory results. To address these limitations, we combined amplicon metabarcoding, microbial isolation and culturing, genomic analysis and molecular microbiology to examine the molecular mechanisms driving cultivar-specific barley colonisation by the rhizobacterial genus *Pseudomonas* [2].

Individual plant species consistently recruit a similar core microbiome that persists regardless of the soil properties, cultivar and even isolation continent [48]. Consistent with this, we observed a substantial degree of overlap between the plant-associated bacterial microbiome samples, with significant enrichment of multiple plant-associated genera, including *Pseudomonas*, compared to bulk soil. Interestingly, the rhizospheres and root-associated microbiota of Chevallier and Tipple barley cultivars also exhibited striking differences, supporting a degree of cultivar-specific microbiome assembly. One such difference was the apparent inability of Tipple to influence the fungal composition of the rhizosphere, in marked contrast to Chevallier, where *Candida* species were almost excluded while *Saitozyma* species were strongly enriched. Focussing on *Pseudomonas*, Chevallier samples displayed a lower abundance of these bacteria than Tipple but higher overall microbial diversity, consistent with the increased complexity of root exudates seen for this landrace.

The differences in *Pseudomonas* recruitment between Chevallier and Tipple were also observed at the level of genotype, with isolated rhizosphere *Pseudomonas* spp. clustering according to their origin. Despite clear evidence for cultivar-dependent genotypic clustering, we observed little overlap between the two biologically independent experiments shown in Figure 2. This is consistent with the underlying soil microbiota exerting a dominant impact on the resulting rhizosphere microbiome [20], but with the barley plants exerting cultivar-specific discrimination on the soil microbiome they find themselves growing in. Bulk soil isolates were evenly distributed throughout the phylogenetic tree, as expected given the soil is the source for most subsequent rhizosphere microbial recruitment.

Our data support a leading role for root exudates in cultivar-specific rhizosphere microbiome assembly in barley. The importance of exudates for root microbe recruitment has been widely studied [33, 49, 50]. Exudates are composed of sugars, amino acids, carboxylic acids, and phenolic compounds, alongside secondary metabolites such as hormones [42], with exudation profiles varying with plant developmental stage, age, plant species and genotype [51]. In barley exudates, the previously uncharacterised flavonoid saponarin has been shown to exert allelopathic properties against weeds such as *Bromus diandrus* [52]. However, comparatively little is known about the relationship between barley root exudates and microbiome assembly.

GC-MS analysis identified several significant differences in the abundance of numerous metabolites between the Chevallier and Tipple exudates, with Tipple exudates containing a less diverse array of compounds overall than Chevallier. We hypothesised that differences in the exudation of key molecules might impact the subsequent establishment of specific microbial communities. The reduced molecular diversity and higher prevalence of hexose sugars, such as glucose and fructose seen in Tipple exudates might therefore explain the greater abundance, but lower phenotypic and genetic diversity observed for the Tipple-associated *Pseudomonas* isolates. In support of this, a *P. fluorescens* mutant (Δ*rccR*) that cannot regulate the glyoxylate shunt and struggles to grow on hexose sugars [44] was significantly compromised in the Tipple rhizosphere, but much less so in Chevallier. The capacity to utilise available carbon resources is clearly a key factor in bacterial adaptation to the rhizosphere environment and is under selection in our experiments. *Pseudomonas* strains that grew well on fructose/glucose were more abundant among the Tipple rhizosphere isolates than isolates from Chevallier.

Analysis of the genomic features present in plant-associated bacterial populations is a powerful tool to identify genes that play important roles in adaptation to plant environments [24, 53]. By examining the frequency of previously identified plant-association loci [45] in the genomes of sequenced barley isolates, we identified a series of genes whose abundance differed significantly between Chevallier and Tipple-derived isolates. A subset of these loci was then deleted in *P. fluorescens* SBW25 and their contribution to *in planta* fitness compared for the two cultivars. Using a single, well-characterised model organism for these assays enabled us to directly compare the impact of the tested genes with one another. Using this approach we identified three new barley cultivar-discrimination genes: *PFLU2583*, a sugar transporter, *PFLU6072*, a transcription factor predicted to play a role in carbon metabolism control; and *PFLU5080*, an uncharacterised protein with a putative role in prophage excision. Differential selection of *PFLU6072* and *PFLU2583* aligns well with the hypothesis that carbon metabolic adaptation is central to successful plant colonisation [44, 54]. The importance of *PFLU5080* is less obvious, although phage-mediated regulation of host behaviour has been identified in several microbes, such as *E. coli* where phage-infected bacteria were shown to outcompete non-infected bacteria under limiting carbon conditions [55]. *PFLU5080* may play an analogous role here, modulating bacterial host responses to environmental differences.

While we identified several cultivar-specific *Pseudomonas* colonisation genes, five of the eight mutants we tested showed little/no fitness defect compared to WT SBW25. This could be due to several reasons, such as differences between our gnotobiotic experimental setup and the original selection environment, genetic differences between SBW25 and the genotypes where these loci were selected, or a degree of functional redundancy in SBW25. This was not entirely unexpected: in a similar study of regulatory genes with potential roles in SBW25 wheat colonisation, only two of seven mutants displayed clear fitness defects, despite all seven showing altered colonisation phenotypes and increased expression in the wheat rhizosphere, consistent with *bona-fide* roles in rhizosphere colonisation [54, 56].

RNA-seq analysis of SBW25 rhizosphere gene expression identified additional loci whose importance differed between Tipple and Chevallier. In addition to around 1000 plant-induced genes whose mRNA abundance increased in both rhizospheres, we identified and tested a subset of cultivar-specific genes. Interestingly, only 14 loci were differently expressed between the two cultivars, with 9 upregulated in the Tipple rhizosphere and 5 in Chevallier. *PFLU3091*, an amino acid permease upregulated in the Tipple rhizosphere and *PFLU4463,* a drug/metabolite transporter upregulated in Chevallier were subsequently shown to be differentially important for colonisation, in agreement with the expression data. These findings demonstrate that barley genotype exerts selective pressure on the *Pseudomonas* population at every organisational level, affecting overall abundance, the frequency of genotypes and individual genetic features and the expression of specific loci within individual microbes.

Previous studies have shown that plants secrete specialised secondary metabolites, such as coumarins [32] or triterpenes [33] to influence their microbiome composition. Our results suggest that higher-level mechanisms for microbial selection also exist, where broad differences in the secretion of primary metabolic compounds can shape the microbiome towards the interests of the plant. The extent to which these two processes for microbiome recruitment co-exist is currently unclear, although it seems plausible that the specialised metabolic shaping described by the Pieterse, Osbourn and Bai labs [32, 33] might act to refine the more general influence of primary exudate secretion we describe here.

While the plant-beneficial properties of rhizosphere microbes are well known [13], the reasons why plant microbiomes differ between members of the same species are less clear. Why do different barley varieties, growing in the same soil environment, recruit different populations of beneficial rhizobacteria and fungi? Excitingly, our results support a cultivar-dependent link between plant growth and the composition of the recruited microbiome, suggesting that the microbiome shaping we see here has real consequences for plant health: plants recruit specific microbes because these microbes help them to grow. Our data suggest that the degree of population fine-tuning that takes place and the consequences of this for plant fitness, are both broader and more complex than we previously suspected. Communication between plants and their microbiota is clearly a two-way conversation.

## Materials and methods

### Biological material and growth conditions

Barley cultivars Chevallier and Tipple were used throughout this study. All seeds were surface sterilised prior to use with 70% ethanol for 1 min, 5% sodium hypochlorite for 2 min followed by thorough washing with sterile distilled water. Following sterilisation, seeds were germinated on 1.5% water agar plates in darkness and at room temperature (RT) for 48 h or until germination before further use.

Bacterial strains used in this work are listed in Supplementary Table 3. *Pseudomonas* strains were grown overnight at 28 °C with shaking in Lysogeny Broth (LB) [57], King’s medium B (KB) [58] or M9 minimal media supplemented with carbon sources at a final concentration of 0.4% w/v [57], as stated in the text. *Pseudomonas* growth media was supplemented with antibiotics and other additives as described elsewhere in the text. *Rhizobium leguminosarum* biosensor strains [39] were grown overnight at 28 °C with shaking in tryptone yeast (TY) [59] or universal minimal salts (UMS) [39] medium. UMS was supplemented with 30mM pyruvate and 10mM ammonium chloride unless specified otherwise and solidified with 1.5% agar where appropriate. Antibiotics were added, when necessary, at the following concentrations: streptomycin (500 μg/ml) and tetracycline (2 μg/ml in UMS, 5 μg/ml in TY). *Escherichia coli* strains were grown in LB medium and plates solidified with 1.5% agar at 37 °C. Media was supplemented with 12.5 μg/ml tetracycline where necessary. *Streptomyces venezuela*e ATCC 10712 was grown at 28 °C until sporulation on MYM medium supplemented with 2% agar [60].

### Isolation of root-associated Pseudomonas

Following germination, barley seedlings were transplanted into 9 cm pots containing JIC (John Innes Centre) Cereal Mix (40% Medium Grade Peat, 40% Sterilised Soil, 20% Horticultural Grit, 1.3kg/m³ PG Mix 14-16-18 + TE (trace elements) Base Fertiliser, 1kg/m³ Osmocote Mini 16-8-11 2mg + TE 0.02% B, Wetting Agent, 3kg/m³ Maglime, 300g/m³ Exemptor). Plants were grown for three weeks in a controlled environment room (CER) at 25 °C with a 16 h light cycle and watered twice a week with sterile tap water. The rhizospheres of three independent plants from each cultivar were sampled individually. Flame-sterilised scissors were used to remove the shoots from each plant and excess soil was removed from the root system by energetic shaking before transferring to sterile 50 ml tubes. 30 ml bulk soil control samples were taken from the centre of unplanted pots. Each tube was filled up to 50 ml with sterile phosphate-buffered saline (PBS) and vortexed for 10 min at 4 °C. Dilution series were produced in sterile PBS and plated onto *Pseudomonas* Agar Base, supplemented with CFC (cetrimide/fucidin/cephalosporin, Oxoid, UK) and prepared according to the manufacturer’s instructions. Plates were incubated at 28 °C until visible colony formation. Single colonies were then randomly selected, streaked to single colonies on KB agar plates and cryopreserved for further analysis. 20 isolates were randomly selected per plant, for a total of 120 isolates per cultivar across two biologically independent experiments.

### Phenotypic characterisation of isolated Pseudomonas spp

Barley rhizosphere *Pseudomonas* isolates were phenotyped for several ecologically relevant traits as follows. A high throughput screening protocol was developed in which 500 cm^2^ square plates were used to assess Congo red binding, UV fluorescence and protease activity. 140 mm petri dishes were used to study *S. venezuelae* suppression and motility and 96-multiwell plates were employed to test HCN production. Plates were inoculated using a microplate replicator. Fluorescence emission under UV light (a proxy for siderophore production) was visually assessed after 48 h growth on KB agar [61]. To assess differences in polysaccharide and proteinaceous adhesin production, 0.005% w/v Congo red dye was added to KB agar and differences in colony pigmentation were assessed after 48 h [61]. Protease production was assessed as the ability to visually degrade 1% w/v milk powder added to KB agar plates after 48 h growth. Motility was studied by observing the spreading patterns of colonies grown for 24 h on 0.5% agar KB plates. Cyanogenic bacteria were detected using an adaptation of the method of Castric and Castric [62], with cultures grown in 96 well plates, after [24]. *S. venezuelae* suppression was assessed for *Pseudomonas* overnight cultures spotted onto a lawn of *Streptomyces* spores (200 µl of a 1:25 suspension spread onto 140 mm petri dishes containing Difco™ Nutrient Agar (DNA, Thermo Fisher, USA)). Growth and inhibition of both microbes was assessed 10 days post-inoculation (dpi).

For each assay, ordinal values between 0 and 3 were assigned to each sample, except for protease activity and motility assays where 0 (phenotype absent) and 1 were assigned. Representative phenotypes for each ordinal value are shown in Supplementary Figure 6. Phenotyping assays were conducted at least twice independently. Where disagreements were recorded in the ordinal data, additional repeats were conducted until a firm consensus was reached.

### Illumina® whole genome sequencing and reciprocal BLAST analysis

22 Chevallier and 20 Tipple derived *Pseudomonas* isolates were selected for whole genome sequencing according to three criteria i) phylogenetic distribution to ensure maximum diversity, ii) growth efficiency in different carbon sources and iii) random selection to minimise bias. Genomic DNA (gDNA) was extracted from overnight LB liquid cultures using a GenElute™ Bacterial Genomic DNA Kit (Sigma-Aldrich, USA) following the manufacturer’s instructions. gDNA quality was evaluated using a NanoDrop ND-1000 Spectrophotometer (Thermo Scientific, USA) and quantity measured with a Qubit® 2.0 Fluorometer using high sensitivity buffer (Thermo Scientific, USA). Sample concentration was normalised to 30 ng/µl and 20 µl of each sample was sent for genome sequencing to the Earlham Institute (Norwich Research Park, UK). LITE libraries were prepared, and Illumina short read sequenced using NovaSeq6000 SP with 150 paired-end reads, aiming for 30x coverage [63].

Genome assembly was performed using SPAdes 3.13.1 and default settings [64], then nucleotide sequences were annotated for the presence of 410 genes of interest using BLAST and custom Perl scripts [65]. The library of genes of interest included known plant-induced genes: transporters, biofilm formation regulators, chemotaxis proteins and siderophore pathways [45] as well as prominent substrate transporters and transcriptional regulators [13, 44]. *P. fluorescens* SBW25 [45] was used as a reference genome. Following reciprocal BLAST analysis, genomes were manually examined for the presence or absence of individual genes. Decisions were made based on a combination of sequence identity (> 66%), alignment coverage (> 66%) and whether each match was a reciprocal best hit. Candidate genes for further mutational analyses were selected manually based on significance (Chi-Squared test, p < 0.05).

### 16S/ITS amplicon sequencing

Five Tipple and five Chevallier barley plants were grown for three weeks in a CER at 25 °C with a 16 h light cycle and watered twice a week with sterile tap water, alongside bulk soil (cereal mix) controls. About 5 g of the rhizosphere-root sample was removed from each plant and decanted into sterile 50 ml tubes. Samples were covered with 10-20 ml of 0.1M KH_2_PO_4_ pH 8.0 buffer and incubated for 30 minutes at room temperature with shaking. This first suspension was kept, and roots were transferred into fresh 50 ml tubes. Another 10-20 ml of buffer was added, and this washing process of the roots was repeated a total of three times, every time keeping the supernatant separately (wash 1, wash 2 and wash 3). For the last step, 10-20 ml were added, and tubes were vortexed twice for 30 s, roots were removed and placed in a new tube and this last wash (wash 4) was collected. Wash 1 and wash 2 were pooled and centrifuged for 10 min at 29000 x g. Supernatant was discharged, wash 3 and 4 were combined into this mix and a final centrifugation was carried out. The resulting pellet was considered to represent the total rhizosphere microbiome. Washed roots were assumed to contain only closely associated epiphytes and endophytes [66].

gDNA extraction was performed using a FastDNA™ SPIN Kit for soil (MP Biomedicals, UK) according to the manufacturer’s instructions. gDNA quality was evaluated using the NanoDrop ND-1000 Spectrophotometer (Thermo Scientific, USA) and quantity measured with a Qubit® 2.0 Fluorometer using high sensitivity buffer (Thermo Scientific, USA). The samples were sent for library preparation and sequencing to Novogene Co. Ltd. (Hong Kong, CN) using Illumina NovaSeq6000 with 250 paired-ends, aiming for 50,000 reads/library. The regions targeted were V4 for the bacterial 16S rRNA gene and ITS1-1F for fungal ITS.

Amplicon sequencing data analysis was conducted on the demultiplexed files as described previously [66] with some modifications. Briefly, on the demultiplexed files supplied with primers and sequence adapters already removed, fastp version 0.20.0 [67] was run on 16S as well as ITS data with disabled length filter and trim_poly_g to remove polyG read tails. Following this, data quality was controlled by R-3.6.3 and DADA2 version 1.14.1 according to the workflow version 2 described in [68]. The truncation length for forward reads was set to 200 bp and for reverse reads to 180 bp for 16S and 210 bp for ITS, respectively. For 16S and ITS libraries the following parameters were used: maxN=0, maxEE=c(2, 2) and truncQ=2 and a minimum length of 50 bp. In both cases, forward and reverse reads were merged with default settings. Silva database (silva_nr_v132) was used to classify bacterial reads [69] and UNITE (sh_general_release_dynamic_s_01.12.2017) for fungal reads [70]. Reads without a match in the databases used were removed. Alpha-diversity analysis was based on Shannon index and Observed measure and calculated on pre-normalised data (package “phyloseq,” R-3.6.3, version 1.30.0). Regarding beta-diversity analysis, ASVs with a mean lower than 10^-5^ were ignored for subsequent analysis, and the filtered ASV data was used to calculate Bray-Curtis beta-diversity (R-3.6.3 “vegan” package, version 2.5.6). Statistical analyses were also performed on filtered data by using the package “vegan”, ANOSIM and PERMANOVA: adonis function [71]. For data visualization ggplot2 (version 3.3.0) was used. Finally, DESeq2 analysis [72] was performed in R-3.6.3 using untransformed data (package ‘DESEq2’, version 1.26.0) to test differentially present taxa between the two cultivars.

### Bacterial genetic manipulation

Vectors and primers used in this work are listed in Supplementary Tables 4 and 5. Gene deletions in SBW25 were made by allelic exchange following a two-step homologous recombination process [73]. To summarise, 500-700 bp homologous flanking regions of the target regions were either PCR-amplified and Gibson-assembled into the BamHI site of the suicide vector pTS1 [74] or synthetically designed by Twist Biosciences (Ca, USA). The pTS1 vector is an adaption of pME3087 [75] containing a *sacB* gene which allows counter-selection on sucrose plates.

*P. fluorescens* SBW25 electrocompetent cells were prepared by growing the cells overnight and washing with 300 mM sucrose three times at RT. These cells were then electroporated at 2500 V with 100-300 ng of the gene deletion constructs. Cells were recovered in 3 ml of LB and incubated for 2 h at 28 °C with shaking to enable expression of antibiotic resistance genes. Single crossovers were selected on 12.5 μg/ml tetracycline and re-streaked to obtain individual colonies. These were then grown overnight without selection in 50 ml of LB at 28 °C with shaking. Dilution series of each culture were plated on LB containing 10% sucrose to counter-select bacteria without a second homologous recombination event. Individual sucrose resistant colonies were patched on LB +/− tetracycline to confirm double recombinants and successful gene deletions confirmed by PCR.

Deletion of the *rccR* gene in the SBW25-*lacZ* background was conducted using the pME3087 construct for *rccR* deletion described in [44]. This deletion vector was transformed by electroporation into SBW25-*lacZ,* and single crossovers selected on 12.5 μg/ml tetracycline and re-streaked to single colonies. Cultures from single crossovers were grown overnight in LB, then diluted 1:100 into fresh medium. After 2 hours, 5 μg/ml tetracycline was added to inhibit the growth of cells that had lost the tetracycline cassette. After a further hour of growth, samples were pelleted and re-suspended in fresh LB containing 5 μg/ml tetracycline and 2 mg/ml piperacillin and phosphomycin to kill growing bacteria. Cultures were grown for a further 4–6 hours, washed once with LB and dilution series were plated onto LB agar. Resulting colonies were patched onto LB +/− tetracycline, and Tet-sensitive colonies tested for gene deletion by colony PCR.

### Bacterial growth assays

Assays were conducted in 96-multiwell plates containing 199 µl of M9 medium supplemented with carbon sources as described in the main text. 1 µl of bacterial overnight LB cultures were inoculated into each well using a multichannel pipette, providing an initial OD_600_ of approx. 0.01. Plates were incubated statically at 28 °C for 48 h and bacterial growth was monitored using a microplate spectrophotometer, either SPECTROstar Nano (BMG LABTECH Ltd., UK) or PowerWave (BioTek Instruments, USA). Each experiment was independently repeated at least twice. ***Root colonisation assays***

Germinated seedlings were placed into sterile 50 ml plastic tubes containing washed, medium grain vermiculite and rooting solution (1 mM CaCl_2_·2H_2_O, 100 μM KCl, 800 μM MgSO_4_, 10 μM FeEDTA, 35 μM H_3_BO_3_, 9 μM MnC_l2_·4H_2_O, 0.8 μM ZnCl_2_, 0.5 μM Na_2_MoO_4_·2H_2_O, 0.3 μM CuSO_4_·5H_2_O, 6 mM KNO_3_, 18.4 mM KH_2_PO_4_, and 20 mM Na_2_HPO_4_). Seedlings were grown for 7 days at 25 °C with 16 h light cycle, then inoculated with a 1:1 mix of 1×10^3^ CFU (Colony Forming Units) wild-type (WT) and mutant SBW25 strains. Plants were grown for a further 5 days, after which shoots were removed and 20 ml of PBS was added to each tube and vortexed for 10 min at 4 °C to resuspend bacteria. Dilution series were then plated onto LB supplemented with 100 µg/ml carbenicillin, 0.1 mM IPTG and 50 µg/ml X-Gal. Plates were incubated at 28 °C until blue and white colonies were clearly distinguishable. Colony counting was undertaken, and final blue/white ratios were calculated for all the mutants tested. Eight to ten plants were sampled per condition and experiments were repeated at least twice independently.

### Root exudate screening with Rlv3841_lux biosensors

The method described in [39] was adapted for optimal results in barley plants. Square 100 mm plates were filled in angle with 75 ml Fahraeus agar (FP) [76], a small square section was pierced on top of each plate to allow growth of the barley seedling and the medium was covered with sterile filter paper upon which one seedling was placed. Biosensors were grown on UMS 1.5% agar slopes for 3 days at 28 °C, then washed three times in UMS medium without any additions. 200-300µl of each Rlv3841_lux biosensor [39] suspension adjusted to an OD_600_ of 0.1 was inoculated directly onto the seedling roots. A second filter paper was applied on top of the inoculated seedling and aluminium foil-wrapped plates were placed in a CER at 25 °C with 16 h light cycle until photographed. A NightOWL II LB 983 CCD camera-box (Berthold Technologies GmbH & Co., Germany) was used to take pictures of plates after two, five and seven dpi. ImageJ software [77] was used to analyse the resulting pictures. The total luminescent area present in each plate was evaluated and relative values were recorded and compared.

### Root exudate extraction and GC-MS analysis

A sterile hydroponic system was constructed and used to grow barley plants for up to three weeks [31]. Two 50 ml plastic tubes were connected with muslin fabric and the bottom part was filled with rooting solution. The whole system was autoclaved to ensure sterility. Barley seedlings were transferred into the tubes and grown in a CER at 25 °C with 16 h light cycle for 3 weeks. The system was topped up with filter-sterilised fresh rooting solution once a week. The root exudates extraction was performed as described elsewhere [31]. Plants were removed from the tubes and roots were carefully washed with sterile deionised water. Four plants were transferred into 200 ml of milli-Q water and incubated for 2 h in the same growing conditions. The liquid fractions were then freeze-dried and diluted with 80% methanol to a concentration of 10 µg/µl. Organic solvent was removed by placing samples on a GeneVac (SP Scientific) prior to processing.

Samples were analysed using Agilent GC-MS Single Quad (7890/5977) plus Gerstel MultiPurpose Sampler (MPS, Agilent Technologies, USA) following the Agilent G1676AA Fiehn GC/MS metabolomics workflow. The MPS enables the automated derivatisation of samples and their subsequent processing. Identification of molecules was based on comparison with the metabolic libraries Agilent Fiehn 2013 GC-MS Metabolomics RTL (Agilent Technologies, USA) and the NIST17 Version 2.3 GC Method Retention Index Library. The Agilent MassHunter Workstation package, particularly Qualitative, Unknowns and Mass Profiler Professional software were used for identification, analysis and data visualisation (Agilent Technologies, USA).

### Rhizosphere RNA extraction and RNA-seq analysis

SBW25 RNA was extracted from the rhizospheres of Chevallier and Tipple growing in axenic conditions as described above for the colonisation assays. 7-day old seedlings were inoculated with 1 ml of a PBS-washed *P. fluorescens* SBW25 culture adjusted to OD_600_ = 1.0 and RNA was extracted at 1 and 5 dpi. 5 ml samples of density-adjusted cell culture prior to plant inoculation were pelleted for 10 min at 9000 x g at 4 °C and included as a control. Pellets were flash-frozen and kept for later RNA extraction.

For the rhizosphere samples, 12 plants were combined to provide sufficient RNA. Three biologically independent rhizosphere samples were extracted per time point. Aerial parts of the plants were removed with sterile scissors and 8 ml of PBS and 12 ml of RNA Later were added. The sample tubes containing vermiculite and separated root systems were then vortexed for 10 min at 4 °C to resuspend bacteria and combined to create a single rhizosphere sample. Immediately, samples were filtered through four layers of previously autoclaved muslin cloth placed in a sterile glass funnel and collected into sterile centrifuge bottles. The filtrate was centrifuged at 170 x g at 4 °C, the supernatant was transferred to a new bottle and the centrifugation step was repeated to remove any residual vermiculite. The supernatant was transferred to a new bottle and centrifuged for 10 min 17000 x g and 4 °C. Cell pellets were then flash-frozen for further use. To lyse the cells, pellets were resuspended in 400 µl of 10mM Tris-Cl pH 8.0 and 700 µl of RLT buffer with β-mercaptoethanol and transferred to Matrix B tubes (MP Biomedicals, UK). Samples were lysed with two 90 second pulses in a FastPrep machine (MP Biomedicals, UK) with 90 seconds rest on ice between them. Samples were centrifuged for 15 min at 16000 x g and 4 °C, and RNA extracted from the supernatant with a RNeasy Mini kit (Qiagen, GE). On-column digestion was conducted with a RNAse-Free DNase Set (Qiagen, GE). RNA quality and concentration was checked by a nanodrop spectrophotometer and a Qubit 2.0 Fluorometer with dilution steps conducted where necessary. Samples were subjected to a second DNase treatment with a TURBO DNA-free Kit (Thermo Scientific, USA) and RNA integrity was confirmed on a 1 % agarose gel.

Samples were sent to Novogene Co. Ltd. (Hong Kong, CN) for rRNA depletion, strand-specific library construction and sequencing on an Illumina NovaSeq 6000. Subread [78] was used to align the reads in the fastq files to the SBW25 reference genome. BAM files were sorted and indexed using SAMtools (version 1.8) [79]. Mapping of the reads were counted from the BAM files using the featureCounts program, part of the Subread package, resulting in a table of counts with one row for each gene and one column for each sample. The table of counts was used as the input to the R package edgeR [80] to test differential expression of each gene and a differential log fold changes table was produced.

### Cross-inoculation experiment

The rhizospheres of 3-week-old barley plants grown in Cereal mix as described above were extracted by pulling the plants from the pots and firmly shaking the roots. The rhizosphere inoculum of each barley cultivar consisted of the combined sample from 4 plants. Shoots were removed and up to 5 g of root material were placed in 50 ml Falcon tubes, which were filled with PBS buffer and vortexed for 10 min at 4 °C to resuspend bacteria. Roots were discarded and the remaining PBS was transferred to a larger container. Heavier particles were removed, and the rhizosphere suspension was centrifuged at 2700 x g for 30 minutes at 4 °C, washed with PBS and the process was repeated three times. The final volume was adjusted so that a sufficient volume of inoculum for each plant could be produced. Each seedling was inoculated with 5ml of this rhizosphere suspension.

The *Pseudomonas* synthetic communities (SynComs) used in the cross-inoculation experiment were created by mixing the 60 strains originally isolated from Chevallier and Tipple roots (see Figure 2) in an equal ratio as described elsewhere [46] but with some modifications. Briefly, overnight cultures were adjusted using PBS instead of MgCl_2_ to an OD_600_ of 0.2, then mixed and re-adjusted to a final OD_600_ = 0.2 with PBS. Seedlings were inoculated with 5 ml of each SynCom inoculum.

The whole experiment was performed under controlled conditions at 25 °C with 16 h light cycle in hydroponic systems using vermiculite and rooting solution as described above. Plants were placed in trays separated by treatment and watering took place from the bottom every two or three days, 200 ml during the first week and 500ml for the rest of the experiment. After three weeks, vermiculite was removed from the roots, plants were placed in an oven at 60 °C for up to six days and dry weight was recorded.

## Figure Legends

**Supplementary Table 1.**
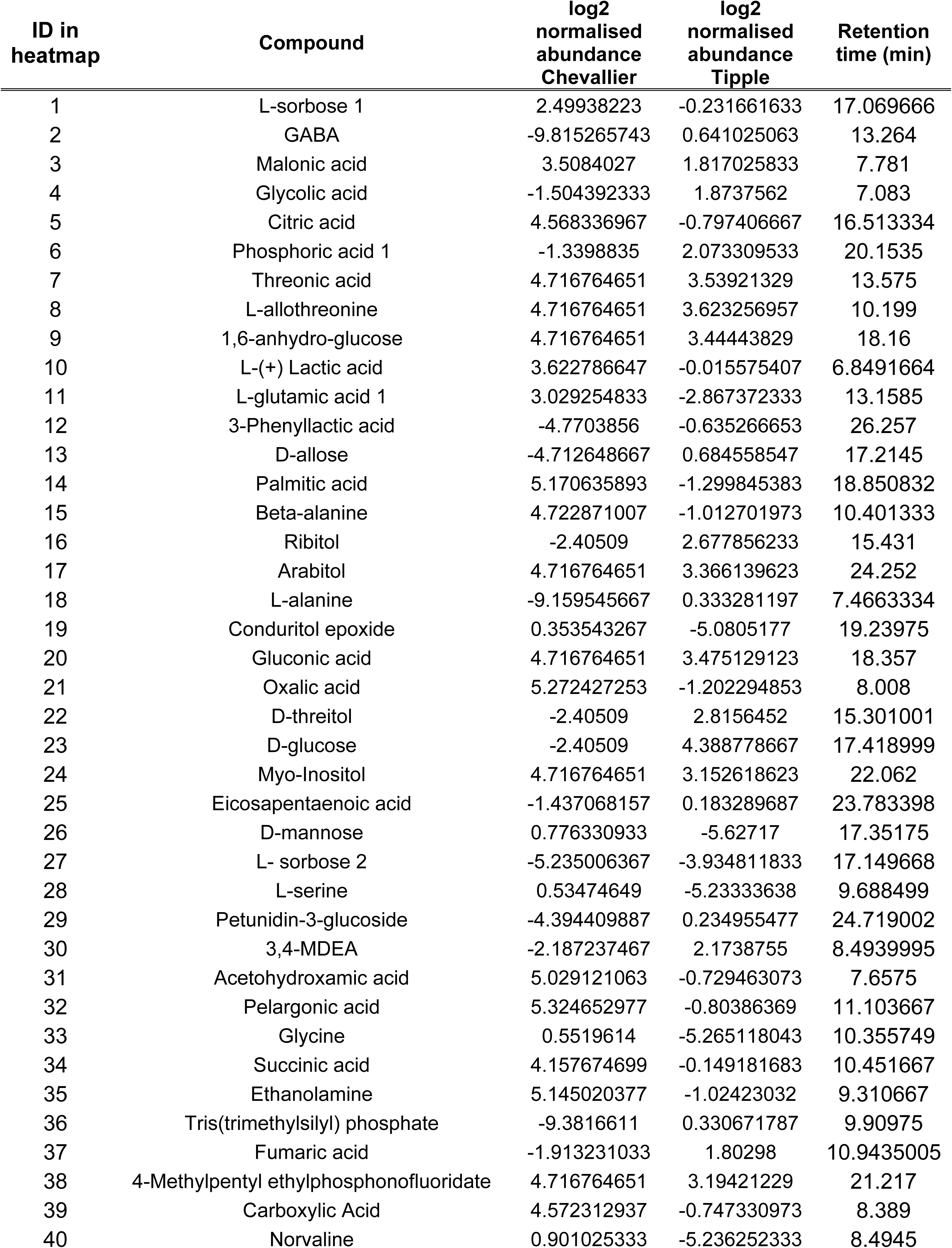
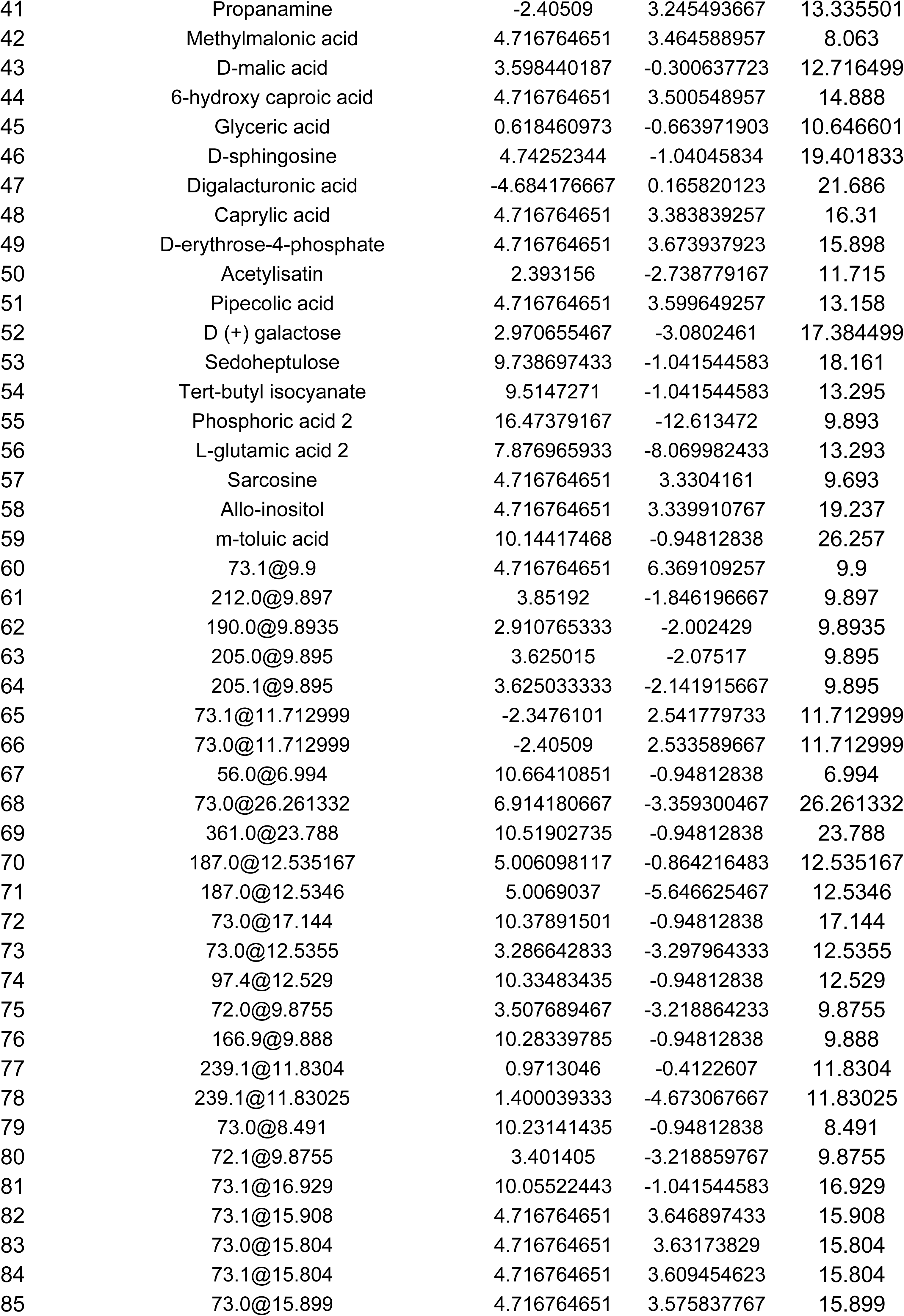
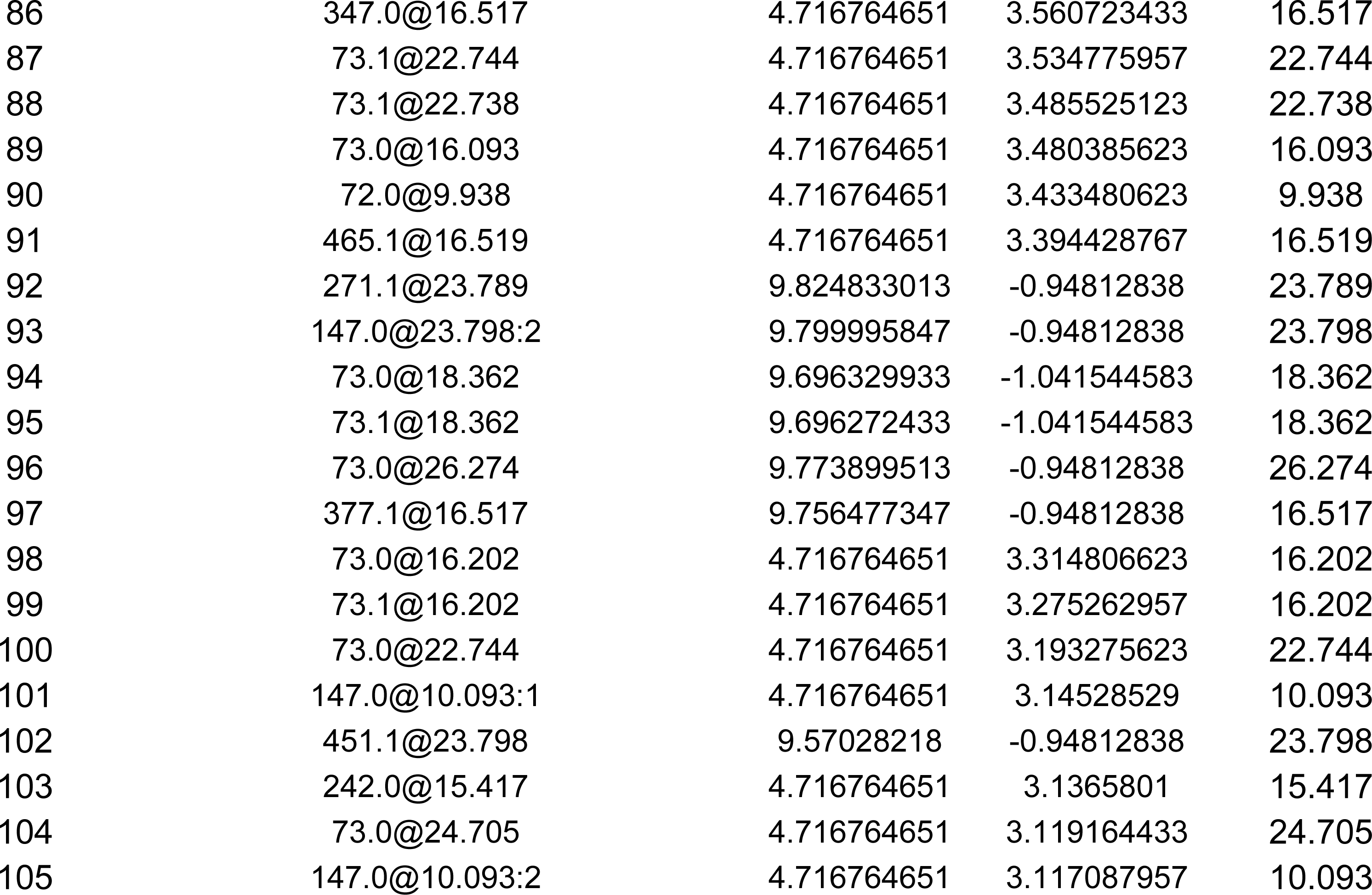
GC-MS detected compounds in root exudates of Tipple and Chevallier.

| ID in heatmap | Compound | log2 normalised abundance Chevallier | log2 normalised abundance Tipple | Retention time (min) |
| --- | --- | --- | --- | --- |
| 1 | L-sorbose 1 | 2.49938223 | -0.231661633 | 17.069666 |
| 2 | GABA | -9.815265743 | 0.641025063 | 13.264 |
| 3 | Malonic acid | 3.5084027 | 1.817025833 | 7.781 |
| 4 | Glycolic acid | -1.504392333 | 1.8737562 | 7.083 |
| 5 | Citric acid | 4.568336967 | -0.797406667 | 16.513334 |
| 6 | Phosphoric acid 1 | -1.3398835 | 2.073309533 | 20.1535 |
| 7 | Threonic acid | 4.716764651 | 3.53921329 | 13.575 |
| 8 | L-allothreonine | 4.716764651 | 3.623256957 | 10.199 |
| 9 | 1,6-anhydro-glucose | 4.716764651 | 3.44443829 | 18.16 |
| 10 | L-(+) Lactic acid | 3.622786647 | -0.015575407 | 6.8491664 |
| 11 | L-glutamic acid 1 | 3.029254833 | -2.867372333 | 13.1585 |
| 12 | 3-Phenyllactic acid | -4.7703856 | -0.635266653 | 26.257 |
| 13 | D-allose | -4.712648667 | 0.684558547 | 17.2145 |
| 14 | Palmitic acid | 5.170635893 | -1.299845383 | 18.850832 |
| 15 | Beta-alanine | 4.722871007 | -1.012701973 | 10.401333 |
| 16 | Ribitol | -2.40509 | 2.677856233 | 15.431 |
| 17 | Arabitol | 4.716764651 | 3.366139623 | 24.252 |
| 18 | L-alanine | -9.159545667 | 0.333281197 | 7.4663334 |
| 19 | Conduritol epoxide | 0.353543267 | -5.0805177 | 19.23975 |
| 20 | Gluconic acid | 4.716764651 | 3.475129123 | 18.357 |
| 21 | Oxalic acid | 5.272427253 | -1.202294853 | 8.008 |
| 22 | D-threitol | -2.40509 | 2.8156452 | 15.301001 |
| 23 | D-glucose | -2.40509 | 4.388778667 | 17.418999 |
| 24 | Myo-Inositol | 4.716764651 | 3.152618623 | 22.062 |
| 25 | Eicosapentaenoic acid | -1.437068157 | 0.183289687 | 23.783398 |
| 26 | D-mannose | 0.776330933 | -5.62717 | 17.35175 |
| 27 | L- sorbose 2 | -5.235006367 | -3.934811833 | 17.149668 |
| 28 | L-serine | 0.53474649 | -5.23333638 | 9.688499 |
| 29 | Petunidin-3-glucoside | -4.394409887 | 0.234955477 | 24.719002 |
| 30 | 3,4-MDEA | -2.187237467 | 2.1738755 | 8.4939995 |
| 31 | Acetohydroxamic acid | 5.029121063 | -0.729463073 | 7.6575 |
| 32 | Pelargonic acid | 5.324652977 | -0.80386369 | 11.103667 |
| 33 | Glycine | 0.5519614 | -5.265118043 | 10.355749 |
| 34 | Succinic acid | 4.157674699 | -0.149181683 | 10.451667 |
| 35 | Ethanolamine | 5.145020377 | -1.02423032 | 9.310667 |
| 36 | Tris(trimethylsilyl) phosphate | -9.3816611 | 0.330671787 | 9.90975 |
| 37 | Fumaric acid | -1.913231033 | 1.80298 | 10.9435005 |
| 38 | 4-Methylpentyl ethylphosphonofluoridate | 4.716764651 | 3.19421229 | 21.217 |
| 39 | Carboxylic Acid | 4.572312937 | -0.747330973 | 8.389 |
| 40 | Norvaline | 0.901025333 | -5.236252333 | 8.4945 |
| 41 | Propanamine | -2.40509 | 3.245493667 | 13.335501 |
| 42 | Methylmalonic acid | 4.716764651 | 3.464588957 | 8.063 |
| 43 | D-malic acid | 3.598440187 | -0.300637723 | 12.716499 |
| 44 | 6-hydroxy caproic acid | 4.716764651 | 3.500548957 | 14.888 |
| 45 | Glyceric acid | 0.618460973 | -0.663971903 | 10.646601 |
| 46 | D-sphingosine | 4.74252344 | -1.04045834 | 19.401833 |
| 47 | Digalacturonic acid | -4.684176667 | 0.165820123 | 21.686 |
| 48 | Caprylic acid | 4.716764651 | 3.383839257 | 16.31 |
| 49 | D-erythrose-4-phosphate | 4.716764651 | 3.673937923 | 15.898 |
| 50 | Acetylisatin | 2.393156 | -2.738779167 | 11.715 |
| 51 | Pipecolic acid | 4.716764651 | 3.599649257 | 13.158 |
| 52 | D (+) galactose | 2.970655467 | -3.0802461 | 17.384499 |
| 53 | Sedoheptulose | 9.738697433 | -1.041544583 | 18.161 |
| 54 | Tert-butyl isocyanate | 9.5147271 | -1.041544583 | 13.295 |
| 55 | Phosphoric acid 2 | 16.47379167 | -12.613472 | 9.893 |
| 56 | L-glutamic acid 2 | 7.876965933 | -8.069982433 | 13.293 |
| 57 | Sarcosine | 4.716764651 | 3.3304161 | 9.693 |
| 58 | Allo-inositol | 4.716764651 | 3.339910767 | 19.237 |
| 59 | m-toluic acid | 10.14417468 | -0.94812838 | 26.257 |
| 60 | 73.1@9.9 | 4.716764651 | 6.369109257 | 9.9 |
| 61 | 212.0@9.897 | 3.85192 | -1.846196667 | 9.897 |
| 62 | 190.0@9.8935 | 2.910765333 | -2.002429 | 9.8935 |
| 63 | 205.0@9.895 | 3.625015 | -2.07517 | 9.895 |
| 64 | 205.1@9.895 | 3.625033333 | -2.141915667 | 9.895 |
| 65 | 73.1@11.712999 | -2.3476101 | 2.541779733 | 11.712999 |
| 66 | 73.0@11.712999 | -2.40509 | 2.533589667 | 11.712999 |
| 67 | 56.0@6.994 | 10.66410851 | -0.94812838 | 6.994 |
| 68 | 73.0@26.261332 | 6.914180667 | -3.359300467 | 26.261332 |
| 69 | 361.0@23.788 | 10.51902735 | -0.94812838 | 23.788 |
| 70 | 187.0@12.535167 | 5.006098117 | -0.864216483 | 12.535167 |
| 71 | 187.0@12.5346 | 5.0069037 | -5.646625467 | 12.5346 |
| 72 | 73.0@17.144 | 10.37891501 | -0.94812838 | 17.144 |
| 73 | 73.0@12.5355 | 3.286642833 | -3.297964333 | 12.5355 |
| 74 | 97.4@12.529 | 10.33483435 | -0.94812838 | 12.529 |
| 75 | 72.0@9.8755 | 3.507689467 | -3.218864233 | 9.8755 |
| 76 | 166.9@9.888 | 10.28339785 | -0.94812838 | 9.888 |
| 77 | 239.1@11.8304 | 0.9713046 | -0.4122607 | 11.8304 |
| 78 | 239.1@11.83025 | 1.400039333 | -4.673067667 | 11.83025 |
| 79 | 73.0@8.491 | 10.23141435 | -0.94812838 | 8.491 |
| 80 | 72.1@9.8755 | 3.401405 | -3.218859767 | 9.8755 |
| 81 | 73.1@16.929 | 10.05522443 | -1.041544583 | 16.929 |
| 82 | 73.1@15.908 | 4.716764651 | 3.646897433 | 15.908 |
| 83 | 73.0@15.804 | 4.716764651 | 3.63173829 | 15.804 |
| 84 | 73.1@15.804 | 4.716764651 | 3.609454623 | 15.804 |
| 85 | 73.0@15.899 | 4.716764651 | 3.575837767 | 15.899 |
| 86 | 347.0@16.517 | 4.716764651 | 3.560723433 | 16.517 |
| 87 | 73.1@22.744 | 4.716764651 | 3.534775957 | 22.744 |
| 88 | 73.1@22.738 | 4.716764651 | 3.485525123 | 22.738 |
| 89 | 73.0@16.093 | 4.716764651 | 3.480385623 | 16.093 |
| 90 | 72.0@9.938 | 4.716764651 | 3.433480623 | 9.938 |
| 91 | 465.1@16.519 | 4.716764651 | 3.394428767 | 16.519 |
| 92 | 271.1@23.789 | 9.824833013 | -0.94812838 | 23.789 |
| 93 | 147.0@23.798:2 | 9.799995847 | -0.94812838 | 23.798 |
| 94 | 73.0@18.362 | 9.696329933 | -1.041544583 | 18.362 |
| 95 | 73.1@18.362 | 9.696272433 | -1.041544583 | 18.362 |
| 96 | 73.0@26.274 | 9.773899513 | -0.94812838 | 26.274 |
| 97 | 377.1@16.517 | 9.756477347 | -0.94812838 | 16.517 |
| 98 | 73.0@16.202 | 4.716764651 | 3.314806623 | 16.202 |
| 99 | 73.1@16.202 | 4.716764651 | 3.275262957 | 16.202 |
| 100 | 73.0@22.744 | 4.716764651 | 3.193275623 | 22.744 |
| 101 | 147.0@10.093:1 | 4.716764651 | 3.14528529 | 10.093 |
| 102 | 451.1@23.798 | 9.57028218 | -0.94812838 | 23.798 |
| 103 | 242.0@15.417 | 4.716764651 | 3.1365801 | 15.417 |
| 104 | 73.0@24.705 | 4.716764651 | 3.119164433 | 24.705 |
| 105 | 147.0@10.093:2 | 4.716764651 | 3.117087957 | 10.093 |

**Supplementary Table 3.**
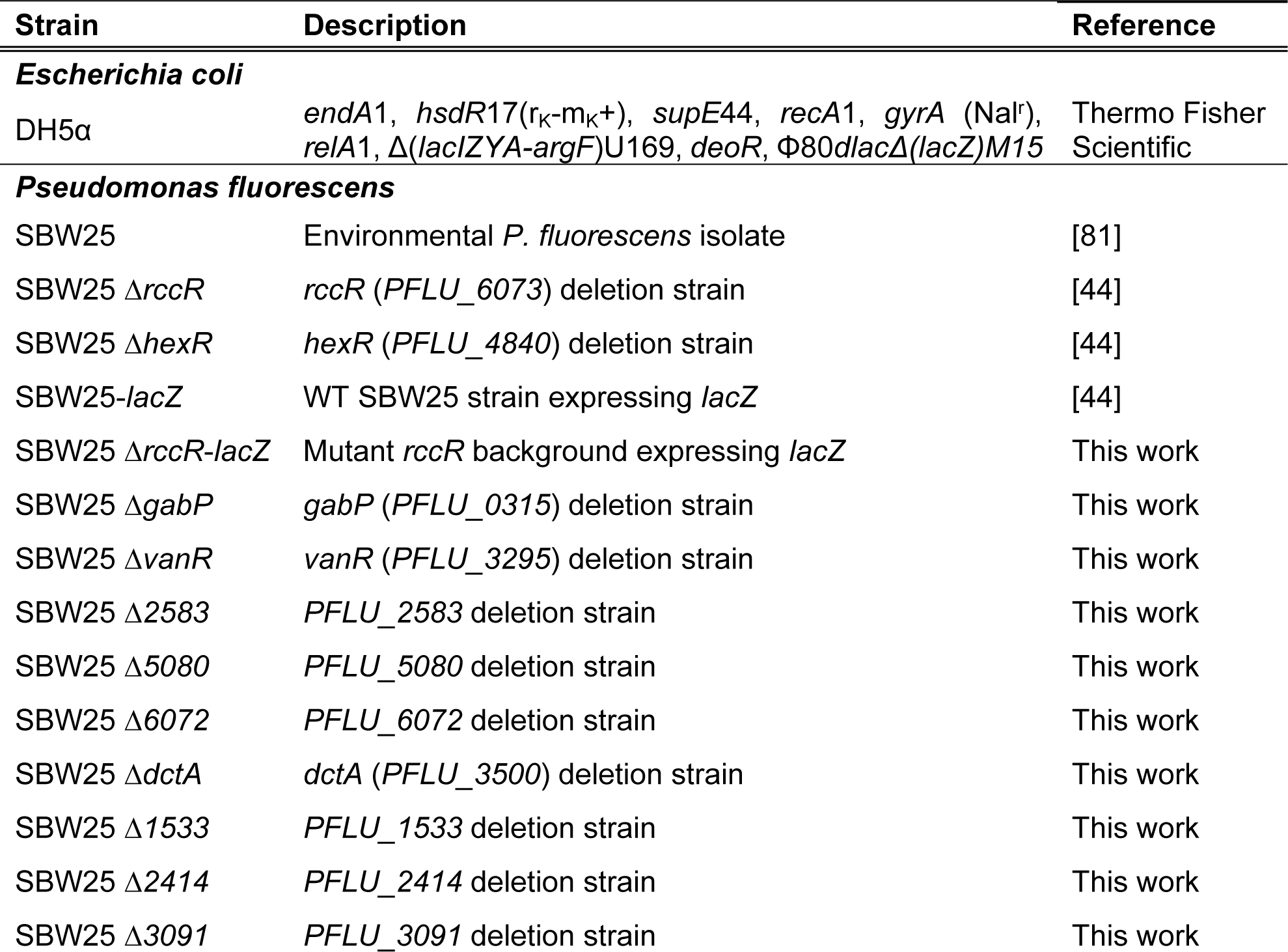
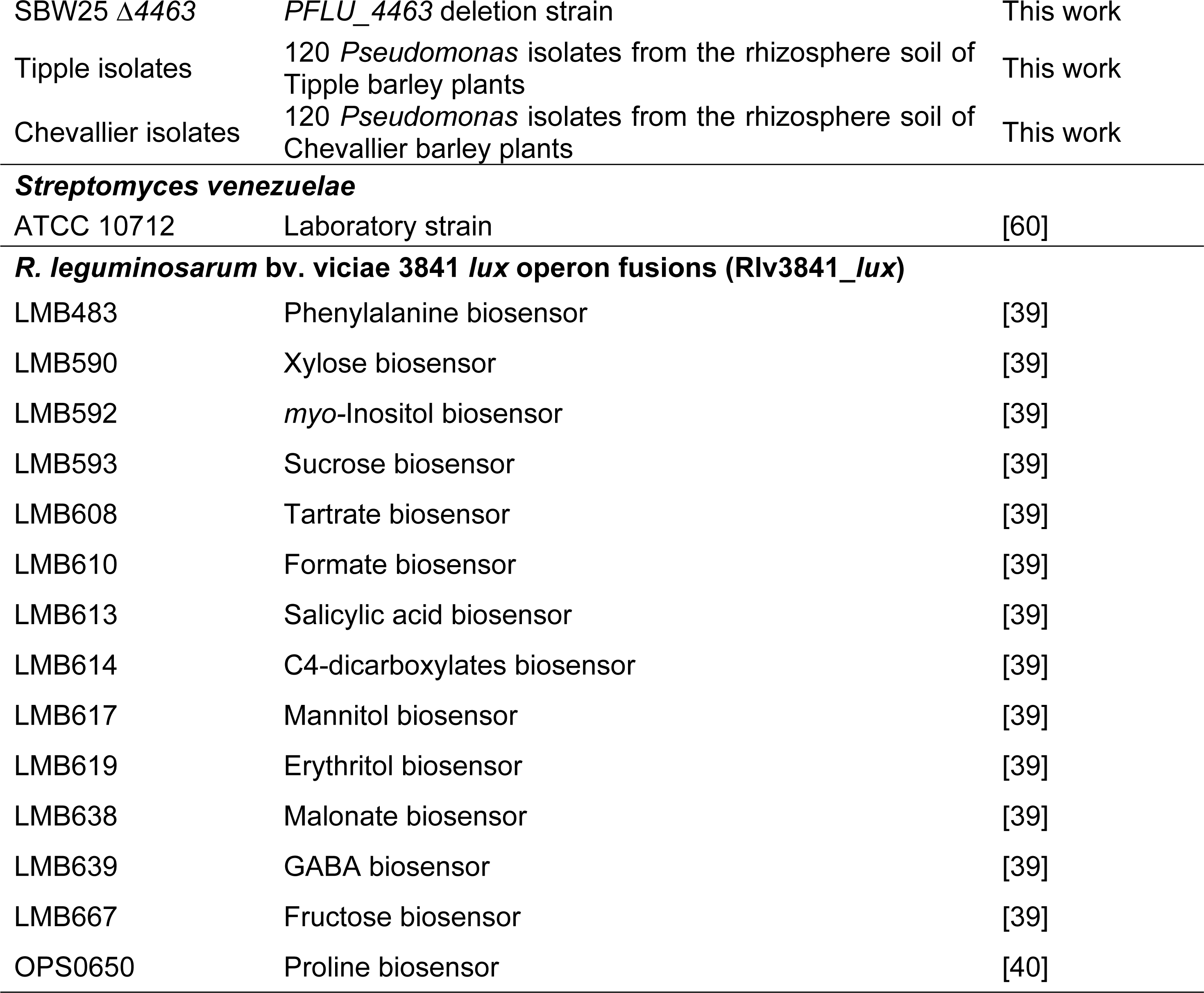
Bacterial strains used in this work.

| Strain | Description | Reference |
| --- | --- | --- |
| ***Escherichia coli*** |  |  |
| DH5α | *endA1, hsdR17(r<sub>K</sub>-m<sub>K</sub><sup>+</sup>), supE44, recA1, gyrA (Nal<sup>r</sup>), relA1, Δ(lacIZYA-argF)U169, deoR, Φ80dlacΔ(lacZ)M15* | Thermo Fisher Scientific |
| ***Pseudomonas fluorescens*** |  |  |
| SBW25 | Environmental *P. fluorescens* isolate | [81] |
| SBW25 Δ*rccR* | *rccR* (PFLU_6073) deletion strain | [44] |
| SBW25 Δ*hexR* | *hexR* (PFLU_4840) deletion strain | [44] |
| SBW25-*lacZ* | WT SBW25 strain expressing *lacZ* | [44] |
| SBW25 Δ*rccR-lacZ* | Mutant *rccR* background expressing *lacZ* | This work |
| SBW25 Δ*gabP* | *gabP* (PFLU_0315) deletion strain | This work |
| SBW25 Δ*vanR* | *vanR* (PFLU_3295) deletion strain | This work |
| SBW25 Δ2583 | PFLU_2583 deletion strain | This work |
| SBW25 Δ5080 | PFLU_5080 deletion strain | This work |
| SBW25 Δ6072 | PFLU_6072 deletion strain | This work |
| SBW25 Δ*dctA* | *dctA* (PFLU_3500) deletion strain | This work |
| SBW25 Δ1533 | PFLU_1533 deletion strain | This work |
| SBW25 Δ2414 | PFLU_2414 deletion strain | This work |
| SBW25 Δ3091 | PFLU_3091 deletion strain | This work |
| SBW25 $\Delta 4463$ | *PFLU_4463* deletion strain | This work |
| Tipple isolates | 120 *Pseudomonas* isolates from the rhizosphere soil of Tipple barley plants | This work |
| Chevallier isolates | 120 *Pseudomonas* isolates from the rhizosphere soil of Chevallier barley plants | This work |
| ***Streptomyces venezuelae*** |  |  |
| ATCC 10712 | Laboratory strain | [60] |
| ***R. leguminosarum* bv. *viciae* 3841 *lux* operon fusions (*Rlv3841\_lux*)** |  |  |
| LMB483 | Phenylalanine biosensor | [39] |
| LMB590 | Xylose biosensor | [39] |
| LMB592 | *myo*-Inositol biosensor | [39] |
| LMB593 | Sucrose biosensor | [39] |
| LMB608 | Tartrate biosensor | [39] |
| LMB610 | Formate biosensor | [39] |
| LMB613 | Salicylic acid biosensor | [39] |
| LMB614 | C4-dicarboxylates biosensor | [39] |
| LMB617 | Mannitol biosensor | [39] |
| LMB619 | Erythritol biosensor | [39] |
| LMB638 | Malonate biosensor | [39] |
| LMB639 | GABA biosensor | [39] |
| LMB667 | Fructose biosensor | [39] |
| OPS0650 | Proline biosensor | [40] |

**Supplementary Table 4.** Plasmids used in this study.

| Plasmid | Description | Antibiotic Resistance | Reference |
| --- | --- | --- | --- |
| pTS1 | Suicide plasmid, pME3087 derivative, *sacB* | Tet | [74] |
| pME3087_$\Delta 6073$ | pME3087 containing 500 bp flanking regions of *PFLU_6073* | Tet | [44] |
| pTS1_$\Delta PFLU0315$ | pTS1 based synthetic construct for deletion of *PFLU_0315* | Tet | This work |
| pTS1_$\Delta PFLU3295$ | pTS1 based synthetic construct for deletion of *PFLU_3295* | Tet | This work |
| pTS1_$\Delta PFLU5080$ | pTS1 based synthetic construct for deletion of *PFLU_5080* | Tet | This work |
| pTS1_$\Delta PFLU6072$ | pTS1 based synthetic construct for deletion of *PFLU_6072* | Tet | This work |
| pTS1_$\Delta PFLU3500$ | pTS1 based synthetic construct for deletion of *PFLU_3500* | Tet | This work |
| pTS1_$\Delta PFLU1533$ | pTS1 based synthetic construct for deletion of *PFLU_1533* | Tet | This work |
| pTS1_$\Delta PFLU2414$ | pTS1 based synthetic construct for deletion of *PFLU_2414* | Tet | This work |

**Supplementary Table 5.**
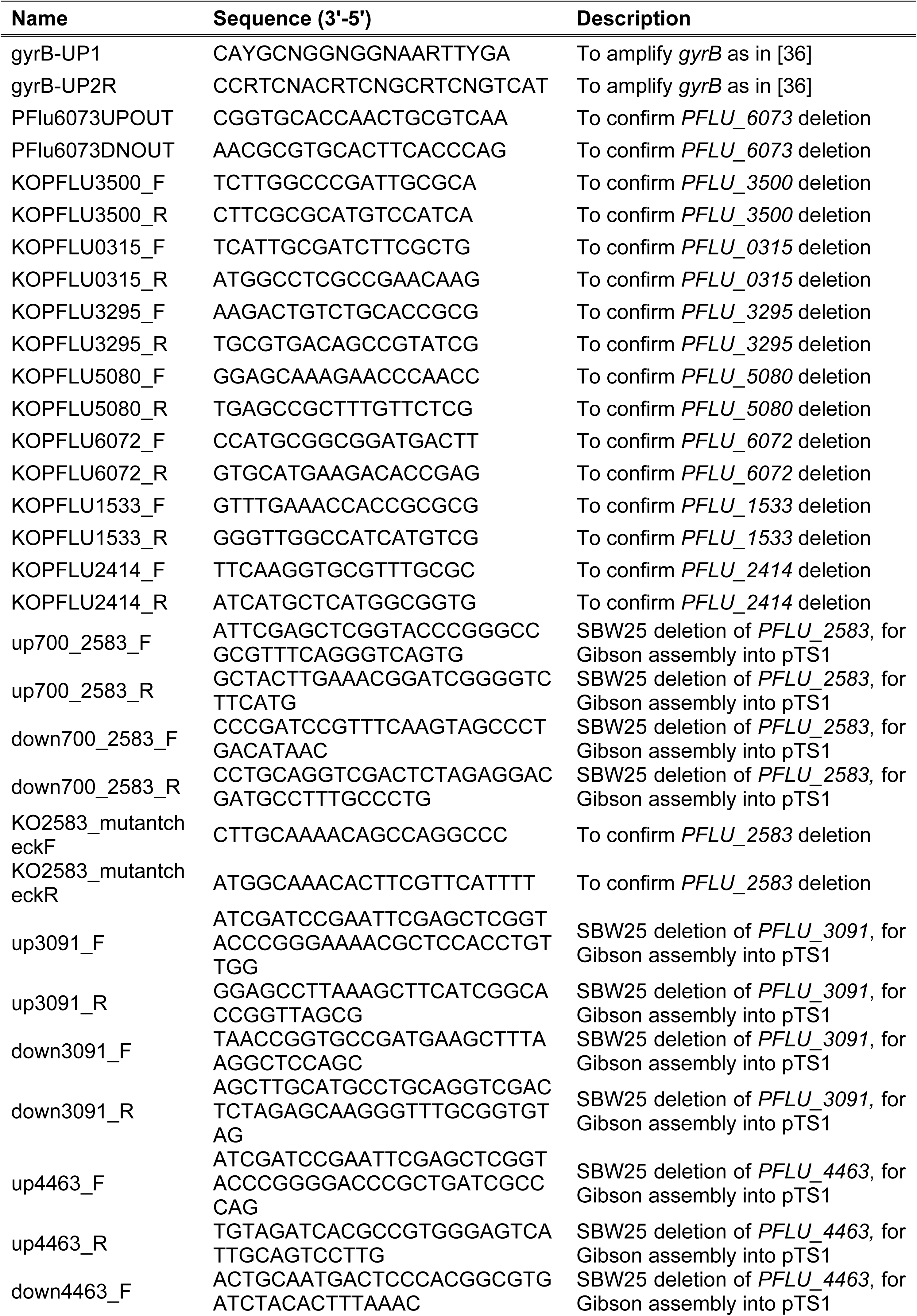
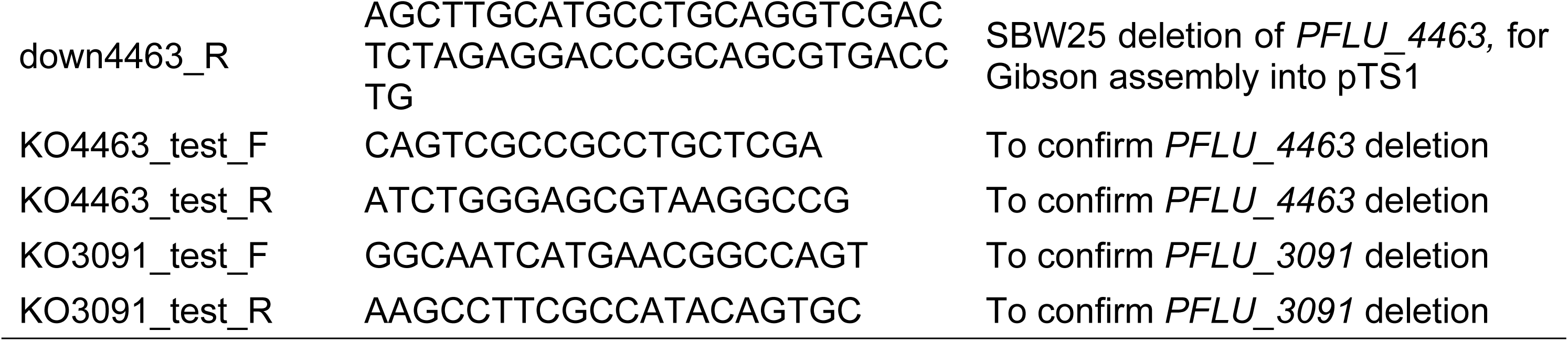
Oligonucleotides used in this work.

**Supplementary figure 1.** Alpha diversity comparison for bacterial communities. Observed richness and Shannon diversity were used as diversity measures. A. Overall comparison between the rhizosphere communities of Chevallier and Tipple and the bulk soil. B. Root endosphere community comparison between Chevallier and Tipple. C. Comparison of community composition between Chevallier compartments. D. Comparison of community composition between Tipple compartments. Five replicates, represented as different coloured dots, were used per condition. Asterisks indicate p < 0.05 (*), 0.01 (**) or 0.001(***).

**Supplementary figure 2.** Alpha diversity comparison for fungal communities. Observed richness and Shannon diversity were used as diversity measures. A. Overall comparison between the rhizosphere communities of Chevallier and Tipple and the bulk soil. Significant differences for both indices, between cultivars and Chevallier and bulk soil. B. Root endosphere community comparison between Chevallier and Tipple. No significant differences were found. C. Comparison of community composition between Tipple compartments. D. Comparison of community composition between Chevallier compartments. Five replicates, represented as different coloured dots, were used per condition. Asterisks indicate p < 0.05 (*), 0.01 (**) or 0.001(***).

**Supplementary figure 3.** Phenotypic traits evaluated in the *Pseudomonas* rhizosphere strains and their significance according to Chi-Square comparisons. A. Congo Red binding (CRB). B. Protease activity (PA). C. Motility (MO). D. CRB Chi-Square paired comparisons. E. PA Chi-Square paired comparisons. F. MO Chi-Square paired comparisons. G. Fluorescence emission (FE). H. Hydrogen cyanide production (HCN). I. *Streptomyces* suppression (SS). J. FE Chi-Square paired comparisons. K. HCN Chi-Square paired comparisons. L. SS Chi-Square paired comparisons. Data is shown as the relative percentage of isolates presenting a given score Significant differences according to Chi-square test are represented.

**Supplementary figure 4.** Growth curves of deletion mutants in SBW25 of genes differentially distributed between Chevallier and Tipple rhizosphere isolates at 36 h. A. LB medium. B. KB medium. C. M9 minimal medium 0.4% acetate. D. M9 minimal medium 0.4% succinate. E. M9 minimal medium 0.4% pyruvate. F. M9 minimal medium 0.4% glucose. G. M9 minimal medium 0.4% Glycerol. 3 biological reps used per strain. Error bars are represented as SEM. Experiment was repeated three times and here a representative graph is shown.

**Supplementary figure 5.** Genomic context of the *PFLU_3089 – PFLU_3098* region of the SBW25 genome. Gene numbers are indicated above each gene. Genes upregulated in the Tipple rhizosphere relative to Chevallier are highlighted in green. Predicted encoded protein functions are given for upregulated genes. Arrows indicate the direction of open reading frame transcription in each case.

**Supplementary figure 6.** Ordinal scales used to score bacterial phenotypes: Congo Red binding (CRB), fluorescence emission (FE), Hydrogen Cyanide production (HCN), Streptomyces suppression (SS), protease activity (PA) and motility (MO).

## References

1. Turner TR, James EK, Poole PS. The plant microbiome. Genome biology. 2013;14(6):209.

2. Muller DB, Vogel C, Bai Y, Vorholt JA. The Plant Microbiota: Systems-Level Insights and Perspectives. Annual review of genetics. 2016;50:211–34. Epub 2016/09/21. doi: 10.1146/annurev-genet-120215-034952. PubMed PMID: 27648643.

3. Orozco-Mosqueda MDC, Rocha-Granados MDC, Glick BR, Santoyo G. Microbiome engineering to improve biocontrol and plant growth-promoting mechanisms. Microbiol Res. 2018;208:25–31. Epub 2018/03/20. doi: 10.1016/j.micres.2018.01.005. PubMed PMID: 29551209.

4. Bulgarelli D, Schlaeppi K, Spaepen S, Ver Loren van Themaat E, Schulze-Lefert P. Structure and functions of the bacterial microbiota of plants. Annual review of plant biology. 2013;64:807–38. Epub 2013/02/05. doi: 10.1146/annurev-arplant-050312-120106. PubMed PMID: 23373698.

5. Tkacz A, Poole P. Role of root microbiota in plant productivity. J Exp Bot. 2015;66(8):2167–75. Epub 2015/04/25. doi: 10.1093/jxb/erv157. PubMed PMID: 25908654; PubMed Central PMCID: PMCPMC4986727.

6. Pfeilmeier S, Petti GC, Bortfeld-Miller M, Daniel B, Field CM, Sunagawa S, et al. The plant NADPH oxidase RBOHD is required for microbiota homeostasis in leaves. Nat Microbiol. 2021;6(7):852–64. Epub 2021/07/02. doi: 10.1038/s41564-021-00929-5. PubMed PMID: 34194036.

7. Hacquard S. Disentangling the factors shaping microbiota composition across the plant holobiont. New Phytologist. 2016;209(2):454–7.

8. Berendsen RL, Pieterse CM, Bakker PA. The rhizosphere microbiome and plant health. Trends in plant science. 2012;17(8):478–86. doi: 10.1016/j.tplants.2012.04.001. PubMed PMID: 22564542.

9. Haichar FeZ, Santaella C, Heulin T, Achouak W. Root exudates mediated interactions belowground. Soil Biology and Biochemistry. 2014;77:69–80. doi: 10.1016/j.soilbio.2014.06.017.

10. Lundberg DS, Lebeis SL, Paredes SH, Yourstone S, Gehring J, Malfatti S, et al. Defining the core Arabidopsis thaliana root microbiome. Nature. 2012;488(7409):86–90. doi: 10.1038/nature11237. PubMed PMID: 22859206; PubMed Central PMCID: PMCPMC4074413.

11. Coats VC, Rumpho ME. The rhizosphere microbiota of plant invaders: an overview of recent advances in the microbiomics of invasive plants. Front Microbiol. 2014;5:368. Epub 2014/08/08. doi: 10.3389/fmicb.2014.00368. PubMed PMID: 25101069; PubMed Central PMCID: PMCPMC4107844.

12. Lugtenberg B, Kamilova F. Plant-growth-promoting rhizobacteria. Annual review of microbiology. 2009;63:541–56.

13. Compant S, Clément C, Sessitsch A. Plant growth-promoting bacteria in the rhizo- and endosphere of plants: Their role, colonization, mechanisms involved and prospects for utilization. Soil Biology and Biochemistry. 2010;42(5):669–78. doi: 10.1016/j.soilbio.2009.11.024.

14. Zboralski A, Filion M. Genetic factors involved in rhizosphere colonization by phytobeneficial Pseudomonas spp. Computational and Structural Biotechnology Journal. 2020.

15. Silby MW, Winstanley C, Godfrey SA, Levy SB, Jackson RW. Pseudomonas genomes: diverse and adaptable. FEMS microbiology reviews. 2011;35(4):652–80.

16. Mauchline TH, Malone JG. Life in earth - the root microbiome to the rescue? Curr Opin Microbiol. 2017;37:23–8. doi: 10.1016/j.mib.2017.03.005. PubMed PMID: 28437662.

17. Seaton SC, Silby MW. Genetics and Functional Genomics of the Pseudomonas fluorescens Group. In al. DCGe, editor. Genomics of Plant-Associated Bacteria. Berlin Heidelberg: Springer-Verlag; 2014. p. 99–125.

18. Garrido-Sanz D, Meier-Kolthoff JP, Goker M, Martin M, Rivilla R, Redondo-Nieto M. Genomic and Genetic Diversity within the Pseudomonas fluorescens Complex. PLoS One. 2016;11(2):e0150183. doi: 10.1371/journal.pone.0150183. PubMed PMID: 26915094; PubMed Central PMCID: PMCPMC4767706.

19. Loper JE, Hassan KA, Mavrodi DV, Davis EW, 2nd, Lim CK, Shaffer BT, et al. Comparative genomics of plant-associated Pseudomonas spp.: insights into diversity and inheritance of traits involved in multitrophic interactions. PLoS Genet. 2012;8(7):e1002784. Epub 2012/07/14. doi: 10.1371/journal.pgen.1002784. PubMed PMID: 22792073; PubMed Central PMCID: PMC3390384.

20. Wei Z, Gu Y, Friman VP, Kowalchuk GA, Xu Y, Shen Q, et al. Initial soil microbiome composition and functioning predetermine future plant health. Sci Adv. 2019;5(9):eaaw0759. Epub 2019/10/04. doi: 10.1126/sciadv.aaw0759. PubMed PMID: 31579818; PubMed Central PMCID: PMCPMC6760924.

21. Wippel K, Tao K, Niu Y, Zgadzaj R, Kiel N, Guan R, et al. Host preference and invasiveness of commensal bacteria in the Lotus and Arabidopsis root microbiota. Nat Microbiol. 2021;6(9):1150–62. Epub 2021/07/28. doi: 10.1038/s41564-021-00941-9. PubMed PMID: 34312531; PubMed Central PMCID: PMCPMC8387241.

22. Meyer KM, Porch R, Muscettola IE, Vasconcelos ALS, Sherman JK, Metcalf CJE, et al. Plant neighborhood shapes diversity and reduces interspecific variation of the phyllosphere microbiome. The ISME journal. 2022;16(5):1376–87. Epub 2022/01/14. doi: 10.1038/s41396-021-01184-6. PubMed PMID: 35022514; PubMed Central PMCID: PMCPMC9038669.

23. Morella NM, Weng FC, Joubert PM, Metcalf CJE, Lindow S, Koskella B. Successive passaging of a plant-associated microbiome reveals robust habitat and host genotype-dependent selection. Proc Natl Acad Sci U S A. 2020;117(2):1148–59. Epub 2019/12/07. doi: 10.1073/pnas.1908600116. PubMed PMID: 31806755; PubMed Central PMCID: PMCPMC6969547.

24. Pacheco-Moreno A, Stefanato FL, Ford JJ, Trippel C, Uszkoreit S, Ferrafiat L, et al. Pan-genome analysis identifies intersecting roles for Pseudomonas specialized metabolites in potato pathogen inhibition. eLife. 2021;10. Epub 2021/11/19. doi: 10.7554/eLife.71900. PubMed PMID: 34792466.

25. Matsumoto H, Fan X, Wang Y, Kusstatscher P, Duan J, Wu S, et al. Bacterial seed endophyte shapes disease resistance in rice. Nat Plants. 2021;7(1):60–72. Epub 2021/01/06. doi: 10.1038/s41477-020-00826-5. PubMed PMID: 33398157.

26. Fagorzi C, Bacci G, Huang R, Cangioli L, Checcucci A, Fini M, et al. Nonadditive Transcriptomic Signatures of Genotype-by-Genotype Interactions during the Initiation of Plant-Rhizobium Symbiosis. Msystems. 2021;6(1):e00974–20.

27. Bulgarelli D, Garrido-Oter R, Munch PC, Weiman A, Droge J, Pan Y, et al. Structure and function of the bacterial root microbiota in wild and domesticated barley. Cell host & microbe. 2015;17(3):392–403. Epub 2015/03/04. doi: 10.1016/j.chom.2015.01.011. PubMed PMID: 25732064; PubMed Central PMCID: PMCPMC4362959.

28. Escudero-Martinez C, Coulter M, Alegria Terrazas R, Foito A, Kapadia R, Pietrangelo L, et al. Identifying plant genes shaping microbiota composition in the barley rhizosphere. Nature communications. 2022;13(1):1–14.

29. Guo H, Nolan TM, Song G, Liu S, Xie Z, Chen J, et al. FERONIA receptor kinase contributes to plant immunity by suppressing jasmonic acid signaling in Arabidopsis thaliana. Current Biology. 2018;28(20):3316–24. e6.

30. Song Y, Wilson AJ, Zhang XC, Thoms D, Sohrabi R, Song S, et al. FERONIA restricts Pseudomonas in the rhizosphere microbiome via regulation of reactive oxygen species. Nat Plants. 2021;7(5):644–54. Epub 2021/05/12. doi: 10.1038/s41477-021-00914-0. PubMed PMID: 33972713.

31. Zhalnina K, Louie KB, Hao Z, Mansoori N, da Rocha UN, Shi S, et al. Dynamic root exudate chemistry and microbial substrate preferences drive patterns in rhizosphere microbial community assembly. Nat Microbiol. 2018;3(4):470–80. Epub 2018/03/21. doi: 10.1038/s41564-018-0129-3. PubMed PMID: 29556109.

32. Stringlis IA, Yu K, Feussner K, de Jonge R, Van Bentum S, Van Verk MC, et al. MYB72-dependent coumarin exudation shapes root microbiome assembly to promote plant health. Proc Natl Acad Sci U S A. 2018;115(22):E5213–E22. Epub 2018/04/25. doi: 10.1073/pnas.1722335115. PubMed PMID: 29686086; PubMed Central PMCID: PMCPMC5984513.

33. Huang AC, Jiang T, Liu YX, Bai YC, Reed J, Qu B, et al. A specialized metabolic network selectively modulates Arabidopsis root microbiota. Science. 2019;364(6440). Epub 2019/05/11. doi: 10.1126/science.aau6389. PubMed PMID: 31073042.

34. Goddard R, de Vos S, Steed A, Muhammed A, Thomas K, Griggs D, et al. Mapping of agronomic traits, disease resistance and malting quality in a wide cross of two-row barley cultivars. PLoS One. 2019;14(7):e0219042. Epub 2019/07/18. doi: 10.1371/journal.pone.0219042. PubMed PMID: 31314759; PubMed Central PMCID: PMCPMC6636724 Malting Ltd provided funding in the form of salary for DG. This does not alter our adherence to PLOS ONE policies on sharing data and materials. There are no patents, products in development or marketed products associated with this research to declare.

35. Callahan BJ, McMurdie PJ, Holmes SP. Exact sequence variants should replace operational taxonomic units in marker-gene data analysis. The ISME journal. 2017;11(12):2639–43.

36. Yamamoto S, Harayama S. PCR amplification and direct sequencing of gyrB genes with universal primers and their application to the detection and taxonomic analysis of Pseudomonas putida strains. Applied and Environmental Microbiology. 1995;61(3):1104–9. doi: doi:10.1128/aem.61.3.1104-1109.1995.

37. Yamamoto S, Harayama S. Phylogenetic relationships of Pseudomonas putida strains deduced from the nucleotide sequences of gyrB, rpoD and 16S rRNA genes. International Journal of Systematic and Evolutionary Microbiology. 1998;48(3):813–9. doi: https://doi.org/10.1099/00207713-48-3-813.

38. Tamura K, Nei M. Estimation of the number of nucleotide substitutions in the control region of mitochondrial DNA in humans and chimpanzees. Molecular biology and evolution. 1993;10(3):512–26.

39. Pini F, East AK, Appia-Ayme C, Tomek J, Karunakaran R, Mendoza-Suarez M, et al. Bacterial Biosensors for in Vivo Spatiotemporal Mapping of Root Secretion. Plant physiology. 2017;174(3):1289–306. Epub 2017/05/13. doi: 10.1104/pp.16.01302. PubMed PMID: 28495892; PubMed Central PMCID: PMCPMC5490882.

40. Rubia MI, Ramachandran VK, Arrese-Igor C, Larrainzar E, Poole PS. A novel biosensor to monitor proline in pea root exudates and nodules under osmotic stress and recovery. Plant and Soil. 2020;452(1):413–22. doi: 10.1007/s11104-020-04577-2.

41. Eilers KG, Lauber CL, Knight R, Fierer N. Shifts in bacterial community structure associated with inputs of low molecular weight carbon compounds to soil. Soil Biology and Biochemistry. 2010;42(6):896–903.

42. Badri DV, Vivanco JM. Regulation and function of root exudates. Plant, cell & environment. 2009;32(6):666–81.

43. Kamilova F, Kravchenko LV, Shaposhnikov AI, Azarova T, Makarova N, Lugtenberg B. Organic acids, sugars, and L-tryptophane in exudates of vegetables growing on stonewool and their effects on activities of rhizosphere bacteria. Molecular Plant-Microbe Interactions. 2006;19(3):250–6.

44. Campilongo R, Fung RKY, Little RH, Grenga L, Trampari E, Pepe S, et al. One ligand, two regulators and three binding sites: How KDPG controls primary carbon metabolism in Pseudomonas. PLoS Genet. 2017;13(6):e1006839. doi: 10.1371/journal.pgen.1006839. PubMed PMID: 28658302; PubMed Central PMCID: PMCPMC5489143.

45. Silby M, Cerdeno-Tarraga A, Vernikos G, Giddens S, Jackson R, Preston G, et al. Genomic and genetic analyses of diversity and plant interactions of Pseudomonas fluorescens. Genome Biology. 2009;10(5). doi: 10.1186/gb-2009-10-5-r51. PubMed PMID: WOS:000267604200010.

46. Bodenhausen N, Bortfeld-Miller M, Ackermann M, Vorholt JA. A synthetic community approach reveals plant genotypes affecting the phyllosphere microbiota. PLoS Genet. 2014;10(4):e1004283. Epub 2014/04/20. doi: 10.1371/journal.pgen.1004283. PubMed PMID: 24743269; PubMed Central PMCID: PMCPMC3990490.

47. O’Neal L, Akhter S, Alexandre G. A PilZ-Containing Chemotaxis Receptor Mediates Oxygen and Wheat Root Sensing in Azospirillum brasilense. Front Microbiol. 2019;10:312. Epub 2019/03/19. doi: 10.3389/fmicb.2019.00312. PubMed PMID: 30881352; PubMed Central PMCID: PMCPMC6406031.

48. Stopnisek N, Shade A. Persistent microbiome members in the common bean rhizosphere: an integrated analysis of space, time, and plant genotype. The ISME journal. 2021:1–15.

49. Chaparro JM, Badri DV, Bakker MG, Sugiyama A, Manter DK, Vivanco JM. Root exudation of phytochemicals in Arabidopsis follows specific patterns that are developmentally programmed and correlate with soil microbial functions. PloS one. 2013;8(2):e55731.

50. Haichar F, Marol C, Berge O, Rangel-Castro J, Prosser J, Balesdent J, et al. Plant host habitat and root exudates shape soil bacterial community structure. The ISME Journal 2: 1221–1230. https://doi.org/101038/ismej.2008.

51. Vives-Peris V, de Ollas C, Gomez-Cadenas A, Perez-Clemente RM. Root exudates: from plant to rhizosphere and beyond. Plant Cell Rep. 2020;39(1):3–17. Epub 2019/07/28. doi: 10.1007/s00299-019-02447-5. PubMed PMID: 31346716.

52. Bouhaouel I, Richard G, Fauconnier M-L, Ongena M, Franzil L, Gfeller A, et al. Identification of barley (Hordeum vulgare L. subsp. vulgare) root exudates allelochemicals, their autoallelopathic activity and against Bromus diandrus Roth. Germination. Agronomy. 2019;9(7):345.

53. Levy A, Salas Gonzalez I, Mittelviefhaus M, Clingenpeel S, Herrera Paredes S, Miao J, et al. Genomic features of bacterial adaptation to plants. Nat Genet. 2017;50(1):138–50. Epub 2017/12/20. doi: 10.1038/s41588-017-0012-9. PubMed PMID: 29255260; PubMed Central PMCID: PMCPMC5957079.

54. Little RH, Woodcock SD, Campilongo R, Fung RKY, Heal R, Humphries L, et al. Differential Regulation of Genes for Cyclic-di-GMP Metabolism Orchestrates Adaptive Changes During Rhizosphere Colonization by Pseudomonas fluorescens. Frontiers in Microbiology. 2019;10. doi: ARTN 1089 10.3389/fmicb.2019.01089. PubMed PMID: WOS:000468022400001.

55. Argov T, Azulay G, Pasechnek A, Stadnyuk O, Ran-Sapir S, Borovok I, et al. Temperate bacteriophages as regulators of host behavior. Current opinion in microbiology. 2017;38:81–7.

56. Thompson CM, Malone JG. Nucleotide second messengers in bacterial decision making. Curr Opin Microbiol. 2020;55:34–9. Epub 2020/03/17. doi: 10.1016/j.mib.2020.02.006. PubMed PMID: 32172083; PubMed Central PMCID: PMCPMC7322531.

57. Miller JH. Experiments in molecular genetics. 1972.

58. King EO, Ward MK, Raney DE. Two simple media for the demonstration of pyocyanin and fluorescin. The Journal of laboratory and clinical medicine. 1954;44(2):301–7. Epub 1954/08/01. PubMed PMID: 13184240.

59. Beringer J. R factor transfer in Rhizobium leguminosarum. Microbiology. 1974;84(1):188–98.

60. Bibb MJ, Domonkos A, Chandra G, Buttner MJ. Expression of the chaplin and rodlin hydrophobic sheath proteins in Streptomyces venezuelae is controlled by sigma(BldN) and a cognate anti-sigma factor, RsbN. Mol Microbiol. 2012;84(6):1033–49. Epub 2012/05/16. doi: 10.1111/j.1365-2958.2012.08070.x. PubMed PMID: 22582857.

61. Mauchline TH, Chedom-Fotso D, Chandra G, Samuels T, Greenaway N, Backhaus A, et al. An analysis of Pseudomonas genomic diversity in take-all infected wheat fields reveals the lasting impact of wheat cultivars on the soil microbiota. Environ Microbiol. 2015;17(11):4764–78. doi: 10.1111/1462-2920.13038. PubMed PMID: 26337499.

62. Castric KF, Castric PA. Method for rapid detection of cyanogenic bacteria. Appl Environ Microbiol. 1983;45(2):701–2. Epub 1983/02/01. PubMed PMID: 16346217; PubMed Central PMCID: PMCPMC242347.

63. Perez-Sepulveda BM, Heavens D, Pulford CV, Predeus AV, Low R, Webster H, et al. An accessible, efficient and global approach for the large-scale sequencing of bacterial genomes. Genome Biology. 2021;22(1). doi: ARTN 349 10.1186/s13059-021-02536-3. PubMed PMID: WOS:000732962600001.

64. Prjibelski A, Antipov D, Meleshko D, Lapidus A, Korobeynikov A. Using SPAdes De Novo Assembler. Curr Protoc Bioinformatics. 2020;70(1):e102. Epub 2020/06/20. doi: 10.1002/cpbi.102. PubMed PMID: 32559359.

65. Stajich JE, Block D, Boulez K, Brenner SE, Chervitz SA, Dagdigian C, et al. The Bioperl toolkit: Perl modules for the life sciences. Genome Res. 2002;12(10):1611–8. Epub 2002/10/09. doi: 10.1101/gr.361602. PubMed PMID: 12368254; PubMed Central PMCID: PMCPMC187536.

66. Bollmann-Giolai A, Giolai M, Heavens D, Macaulay I, Malone J, Clark MD. A low-cost pipeline for soil microbiome profiling. MicrobiologyOpen. 2020;9(12):e1133.

67. Chen S, Zhou Y, Chen Y, Gu J. fastp: an ultra-fast all-in-one FASTQ preprocessor. Bioinformatics. 2018;34(17):i884–i90.

68. Callahan BJ, Sankaran K, Fukuyama JA, McMurdie PJ, Holmes SP. Bioconductor workflow for microbiome data analysis: from raw reads to community analyses. F1000Research. 2016;5.

69. Quast C, Pruesse E, Yilmaz P, Gerken J, Schweer T, Yarza P, et al. The SILVA ribosomal RNA gene database project: improved data processing and web-based tools. Nucleic acids research. 2012;41(D1):D590–D6.

70. Nilsson RH, Larsson K-H, Taylor AFS, Bengtsson-Palme J, Jeppesen TS, Schigel D, et al. The UNITE database for molecular identification of fungi: handling dark taxa and parallel taxonomic classifications. Nucleic acids research. 2019;47(D1):D259–D64.

71. Dixon P. VEGAN, a package of R functions for community ecology. Journal of Vegetation Science. 2003;14(6):927–30.

72. Love MI, Huber W, Anders S. Moderated estimation of fold change and dispersion for RNA-seq data with DESeq2. Genome biology. 2014;15(12):1–21.

73. Hmelo LR, Borlee BR, Almblad H, Love ME, Randall TE, Tseng BS, et al. Precision-engineering the Pseudomonas aeruginosa genome with two-step allelic exchange. Nature protocols. 2015;10(11):1820–41.

74. Scott TA, Heine D, Qin Z, Wilkinson B. An L-threonine transaldolase is required for L-threo-β-hydroxy-α-amino acid assembly during obafluorin biosynthesis. Nature communications. 2017;8(1):1–11.

75. Voisard C, Bull CT, Keel C, Laville J, Maurhofer M, Schnider U, et al. Biocontrol of root diseases by Pseudomonas fluorescens CHA0: current concepts and experimental approaches. Molecular ecology of rhizosphere microorganisms: biotechnology and the release of GMOs. 1994:67–89.

76. Somasegaran P, Hoben H. Handbook for rhizobia: methods in legume-Rhizobium technology. Handbook for rhizobia: methods in legume-Rhizobium technology. 1994.

77. Rueden CT, Schindelin J, Hiner MC, DeZonia BE, Walter AE, Arena ET, et al. ImageJ2: ImageJ for the next generation of scientific image data. BMC bioinformatics. 2017;18(1):529.

78. Liao Y, Smyth GK, Shi W. The Subread aligner: fast, accurate and scalable read mapping by seed-and-vote. Nucleic Acids Res. 2013;41(10):e108. Epub 2013/04/06. doi: 10.1093/nar/gkt214. PubMed PMID: 23558742; PubMed Central PMCID: PMCPMC3664803.

79. Li H, Handsaker B, Wysoker A, Fennell T, Ruan J, Homer N, et al. The Sequence Alignment/Map format and SAMtools. Bioinformatics. 2009;25(16):2078–9. Epub 2009/06/10. doi: 10.1093/bioinformatics/btp352. PubMed PMID: 19505943; PubMed Central PMCID: PMCPMC2723002.

80. Chen Y, Lun AT, Smyth GK. From reads to genes to pathways: differential expression analysis of RNA-Seq experiments using Rsubread and the edgeR quasi-likelihood pipeline. F1000Res. 2016;5:1438. Epub 2016/08/11. doi: 10.12688/f1000research.8987.2. PubMed PMID: 27508061; PubMed Central PMCID: PMCPMC4934518.

81. Rainey PB, Bailey MJ. Physical and genetic map of the Pseudomonas fluorescens SBW25 chromosome. Mol Microbiol. 1996;19(3):521–33. Epub 1996/02/01. PubMed PMID: 8830243.

